# LF4/MOK and a CDK-related kinase regulate the number and length of cilia in *Tetrahymena*

**DOI:** 10.1101/582833

**Authors:** Yu-Yang Jiang, Wolfgang Maier, Ralf Baumeister, Gregory Minevich, Ewa Joachimiak, Dorota Wloga, Zheng Ruan, Natarajan Kannan, Stephen Bocarro, Anoosh Bahraini, Krishna Kumar Vasudevan, Karl Lechtreck, Eduardo Orias, Jacek Gaertig

## Abstract

The length of cilia is controlled by a poorly understood mechanism that involves members of the conserved RCK kinase group, and among them, the LF4/MOK kinases. In *Tetrahymena*, a loss of an LF4/MOK ortholog, LF4A, lengthened the locomotory cilia, but also reduced their total number per cell. Without LF4A, cilia assembled faster and showed signs of increased intraflagellar transport (IFT). Consistently, overproduced LF4A shortened cilia and downregulated the IFT. GFP-tagged LF4A, expressed in the native locus and imaged by total internal reflection microscopy, was enriched at the basal bodies and distributed along the shafts of cilia. Within cilia, most LF4A-GFP particles were immobile and a few either diffused or moved by IFT. A forward genetic screen identified a CDK-related kinase, CDKR1, whose loss-of-function suppressed the shortening of cilia caused by overexpression of LF4A, by reducing its kinase activity. A loss of CDKR1 alone lengthened both the locomotory and oral cilia. CDKR1 resembles other known ciliary CDK-related kinases: LF2 of *Chlamydomonas*, mammalian CCRK and DYF-18 of *C. elegans,* in lacking the cyclin-binding motif and acting upstream of RCKs. We propose that the total LF4/MOK activity per cilium is dependent on both its activation by an upstream CDK-related kinase and cilium length. Previous studies showed that the rate of assembly is high in growing cilia and decreases as cilia elongate to achieve the steady-state length. We propose that in a longer cilium, the IFT components, which travel from the base to the tip, are subjected to a higher dose of inhibition by the uniformly distributed LF4/MOK. Thus, in a feedback loop, LF4/MOK may translate cilium length into proportional inhibition of IFT, to balance the rates of assembly and disassembly at steady-state.

**Author summary:** Cilia are conserved organelles that generate motility and mediate vital sensory functions, including olfaction and vision. Cilia that are either too short or too long fail to generate proper forces or responses to extracellular signals. Several cilia-based diseases (ciliopathies) are associated with defects in cilia length. Here we use the multiciliated model protist *Tetrahymena,* to study a conserved protein kinase whose activity shortens cilia, LF4/MOK. We find that cells lacking an LF4/MOK kinase of *Tetrahymena*, LF4A, have excessively long, but also fewer cilia. We show that LF4A decreases the intraflagellar transport, a motility that shuttles ciliary precursors from the cilium base to the tip. Live imaging revealed that LF4A is distributed along cilium length and remains mostly immobile, likely due to its anchoring to ciliary microtubules. We proposed that in longer cilia, the intraflagellar transport machinery is exposed to a higher dose of inhibition by LF4A, which could decrease the rate of cilium assembly, to balance the rate of cilium disassembly in mature cilia that maintain stable length.

## Introduction

The classical “long-zero” experiment in the green flagellate *Chlamydomonas reinhardtii* revealed that the length of cilia is regulated [1]. When one of the two cilia of *Chlamydomonas* was removed, the intact cilium immediately started to shorten, while the amputated cilium started to regrow. When both cilia reached about the same intermediate length, they continued to elongate at the same rate, to achieve an equal steady-state length [1]. These observations suggested that cilia length is sensed and actively maintained. A number of ciliopathies including Joubert syndrome [2], Meckel syndrome [3, 4], endocrine-cerebro-osteodysplasia syndrome [5, 6], short rib polydactyly syndrome [7, 8], retinitis pigmentosa [9, 10], non-syndromic recessive deafness [11], polycystic kidney disease [12, 13] and juvenile epilepsy [14] are caused by mutations in proteins that are affect cilium length.

The assembly of most cilia involves delivery of precursors from the cell body to the ciliary base, followed by their distribution along cilium by the intraflagellar transport (IFT) pathway [15]. During IFT, motor proteins move large protein complexes, IFT trains, that in turn ferry precursors of cilia, including tubulin, along axonemal microtubules [16-18]. Kinesin-2 is the IFT motor that operates in the anterograde direction, from the cilium base to the distal tip, where most of the precursors are incorporated into the axoneme [19, 20]. IFT dynein (dynein-2) returns IFT trains and components that turn-over back to the ciliary base [21-23]. The cilium disassembly pathway involves kinesin-related microtubule end destabilizers [24-28], and protein modifications including glutamylation [29], ubiquitination [30, 31] and phosphorylation (reviewed in [32]). In a mature cilium, its steady-state length results from a balance between the rates of assembly (mediated by IFT) and disassembly [33]. Importantly, IFT changes as a function of cilium assembly status. In *Chlamydomonas*, the IFT train size [34, 35] and cargo load [16, 17, 36] are higher in assembling cilia as compared to steady-state or disassembling cilia. On the other hand, also in *Chlamydomonas*, the disassembly rate increases in cilia that are abnormally long [37]. These observations suggest that there are mechanisms that sense and adjust cilium length act by controlling the rates of cilia assembly (IFT) and disassembly.

Several conserved kinases negatively regulate cilium length (their loss makes cilia longer) including: two subgroups of the ROS cross-hybridizing kinases (RCKs): LF4/MOK [37-40] and DYF-5/ICK/MAK [38, 41-46], CDK-related kinases: LF2/CCRK and DYF-18 [43, 46-48], NRK/NEK [37, 49-52] and LF5/CDKL5 [53]. Consistently, inhibition of protein phosphatases (PP1 and PP2A) shortens cilia [54-57]. On the other hand, CDK5 in mammals [58, 59] and CDPK1 (ortholog of the mammalian CMKII) in *Chlamydomonas* [60], promote cilia assembly as their losses make cilia shorter. CDPK1 promotes cilia assembly by increasing the turnaround of IFT trains at the ciliary tip [60]. However, CDPK1 also decreases the assembly rate by inhibiting the entry of IFT trains into cilium [60].

Recent reports have linked some cilium length-regulating kinases to the anterograde IFT motor, kinesin-2. In *Chlamydomonas,* CDPK1 phosphorylates the tail domain of the motor subunit of kinesin-2 (FLA8) and a non-phosphorylatable substitution at this phosphorylation site (S663A) inhibits association of kinesin-2 with IFT complexes and reduces their entry into cilia [60]. In *C. elegans,* loss-of-function mutations in an RCK DYF-5, and a CDK-related DYF-18, rescue defective ciliogenesis caused by an autoinhibitory mutation in the kinesin-2 motor OSM-3, suggesting that DYF-5 and DYF-18 inhibit kinesin-2 [46].

It remains unclear how cilium length is sensed and translated into proper modulations of the rates of cilium assembly and disassembly. The majority of published studies on the control of cilia length have been done in *Chlamydomonas reinhardtii* that carries two cilia, and in animal cells with single primary cilia (reviewed in [61-63]) and less is known about how cilium length is regulated in multiciliated cells, such as the ciliate *Tetrahymena thermophila* used here.

In *Chlamydomonas*, a loss of LF4 (long flagella protein 4, an ortholog of the mammalian MOK) makes cilia twice as long as in the wild type [39, 64], increases the amount of IFT proteins entering the cilium [35] and increases the rates of both cilium assembly and disassembly [37]. Here we investigate the significance of LF4/MOK in the multiciliated *Tetrahymena*. A single *Tetrahymena* cell carries between 500-1000 cilia, including oral cilia that support phagocytosis, and locomotory cilia that are arranged in ∼20 longitudinal rows (reviewed in [65]). Importantly, in *Tetrahymena,* both the time of assembly and length are dependent on cilium type (oral versus locomotory) and position on the anteroposterior cell axis [50, 66, 67]. We find here that *Tetrahymena* has a single cilia-associated LF4/MOK kinase, LF4A, which negatively regulates the length, and positively regulates the number of locomotory cilia. We use a forward genetic screen to identify a CDK-type kinase, CDKR1, as an activator of LF4/MOK, which regulates the length of both locomotory and oral cilia. We propose that cilium length regulation involving LF4/MOK kinases is two-tiered. First, an upstream CDK-related kinase enhances the kinase specific activity of LF4/MOK. Second, due to the ability of LF4/MOK to distribute along cilium length, the aggregated activity of LF4/MOK per cilium may increase as the cilium gets longer, thereby translating the organelle length into a proportional inhibition of IFT. Our observations also point to a link between cilium length and number in multiciliated cells. Finally, our data suggest that specific cilium length-regulating kinases can be targeted to subsets of cilia in the same multiciliated cell.

## Results

### LF4A both shortens cilia and promotes ciliogenesis in *Tetrahymena*

The CMGC (<u>C</u>DK, <u>M</u>AP, <u>G</u>SK, <u>C</u>DK-like [68]) kinase family contains several conserved subfamilies whose members affect cilium length including: RCKs (subdivided into LF4/MOK and DYF-5/MAK/ICK groups), CDK-related kinases (LF2/CCRK and DYF-18) and LF5/CDKL5 kinases. We performed a phylogenetic analysis of the cilia-associated CMGC kinases of *Tetrahymena thermophila* and several ciliated and nonciliated species. The genome of *Tetrahymena* encodes members of each of the cilia-associated CMGC subfamily except for LF2/CCRK (Fig 1A). However, among the CDK-related kinases, TTHERM_01080590 protein (that we will later rename CDKR1, see below) groups with DYF-18 of *C. elegans*, a cilia-associated kinase that was suggested to be a homolog of LF2/CCRKs [46, 48]. Among the RCKs, *Tetrahymena* has seven DYF-5/MAK/ICK type kinases and two LF4/MOK kinases: LF4A (TTHERM_00058800) and LF4B (TTHERM_00822360). While LF4A acts in cilia, surprisingly, LF4B appears to be expressed only during the sexual process of conjugation, where it likely plays a non-ciliary role (see below and S1 Fig).

**Figure 1.**
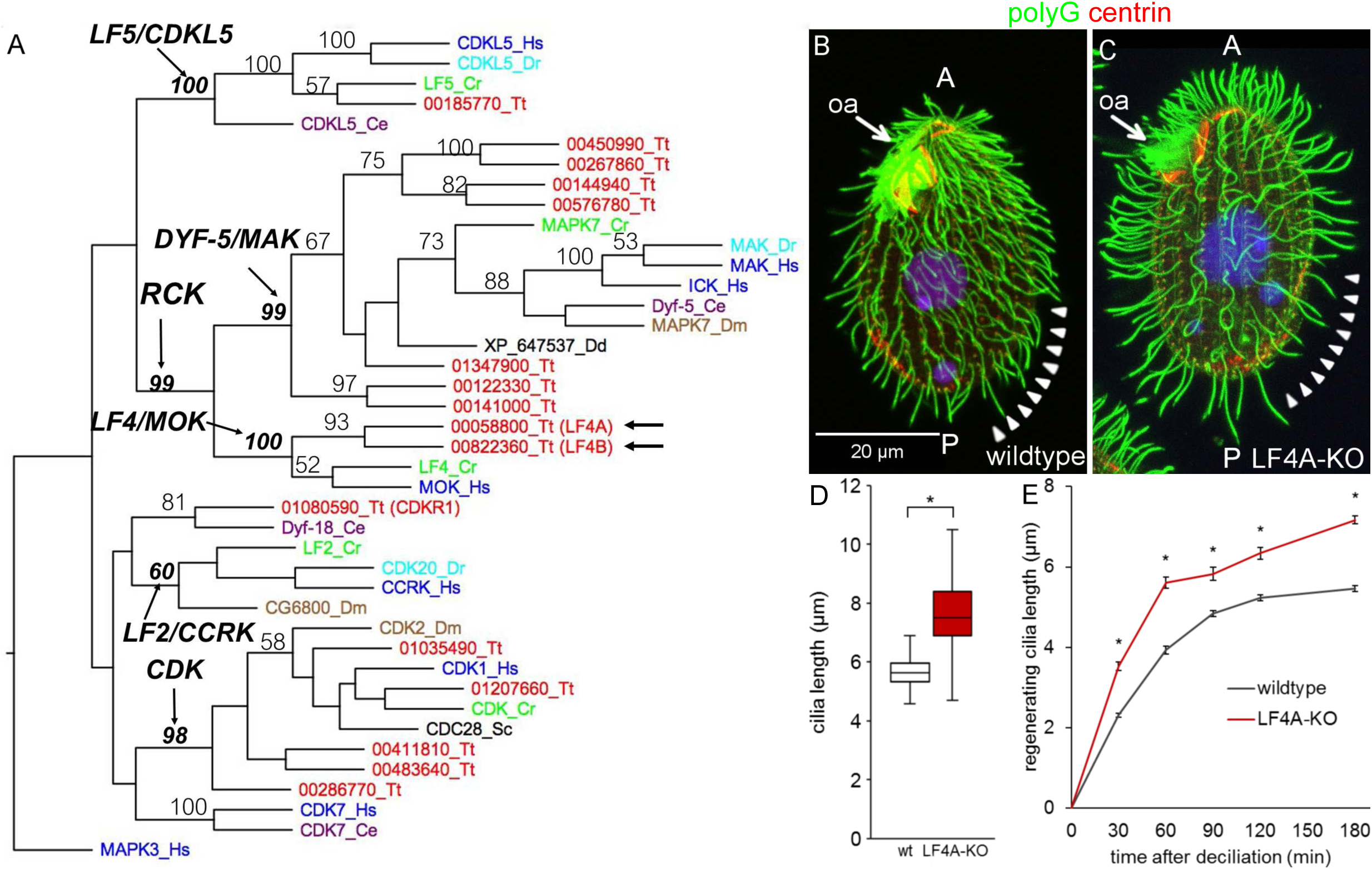
*LF4A* regulates cilia length and number in *Tetrahymena*. (A) A neighbor joining phylogenetic tree of a subset of CMGC kinases. The human MAPK3 was used as an outgroup. The numbers on the branches represent bootstrap support values above 50%. Arrows mark the two LF4/MOK homologs of *Tetrahymena*. (B and C) A wild-type (B) and an LF4A-KO cell (C) stained with the anti-polyG antibodies to visualize cilia (green), anti-centrin antibody 20H5 to mark the basal bodies (red) and DAPI (blue). Arrowheads indicate the posterior-dorsal region that is less densely ciliated in LF4A-KO. Abbreviations: oa, oral apparatus; A, anterior cell end; P, posterior cell end. (D) (D) A box and whisker plot of locomotory cilia length. Cilia from ten wildtype and ten LF4A-KO cells were measured. The average cilium length for the wild type (n = 216) is 5.88 µm, SD (standard deviation): 0.38 µm; LF4A-KO (n = 367) 7.74 µm, SD: 1.82 µm. p < 0.01. (E) The length of cilia during regeneration after deciliation by pH shock. Four to 6 cells, 50 - 150 cilia were used at each time point. Asterisks (*) indicate significant differences (p < 0.01 at each indicated time point).

We constructed *Tetrahymena* cells homozygous for a disruption of the *LF4A* gene (LF4A-KO strain). The LF4A-KO cells assembled fewer locomotory cilia (especially in the posterior/dorsal region) that were 32% longer, compared to the wild type (Fig 1B-1D). In the wild type, cilium length gradually increases from the anterior to the posterior cell end ([50] and Fig 1B). Such a length gradient was also apparent in the LF4A-KO cells despite the overall lengthening of cilia (Fig 1C). The length of oral cilia appeared unaffected (Fig 1B and 1C, arrows). In *Chlamydomonas*, LF4 decreases the rate of cilia assembly [35, 37]. In agreement, following deciliation of *Tetrahymena* by a pH shock, the locomotory cilia grew faster in LF4A-KO than in the wild type (Fig 1E).

### Inside cilia, LF4A-GFP is mostly stationary and rarely diffuses or moves as IFT cargo

We added GFP to the C-terminus of LF4A, by engineering its gene (to preserved expression under the native promoter). Based on immunofluorescence, LF4A-GFP was strongly present at the ciliary bases (both oral and locomotory) and a weaker signal was scattered along the shafts of locomotory cilia (Fig 2B compare to 2A). Total internal reflection fluorescence microscopy (TIRFM) confirmed the enrichment of LF4A-GFP at ciliary bases and its weaker but uniform presence in the shafts of locomotory cilia (Fig 2D). In live cells imaged by TIRFM, most of LF4A-GFP particles were immobile and a few either diffused or moved along linear tracks with IFT velocities (in either anterograde or retrograde direction (Fig 2E, arrows)). The mobile LF4A-GFP particles were rare: not more than a few percent of cilia per cell had convincingly mobile LF4A-GFP and even inside these cilia most of LF4A-GFP particles remained immobile (S1 video). This contrasts with the behavior of fluorescently tagged IFT proteins, most of which were mobile under the same imaging conditions (Fig. 2H, 2H’ and 3H). Thus, most of LF4A may be anchored in the cilium. Based on immunofluorescence, the signal of LF4A-GFP in the ciliary shafts (but not at the basal bodies) greatly decreased after extraction with Triton X-100 (Fig 2C compare to 2B), indicating that the suggested anchorage of LF4A is weak.

**Figure 2.**
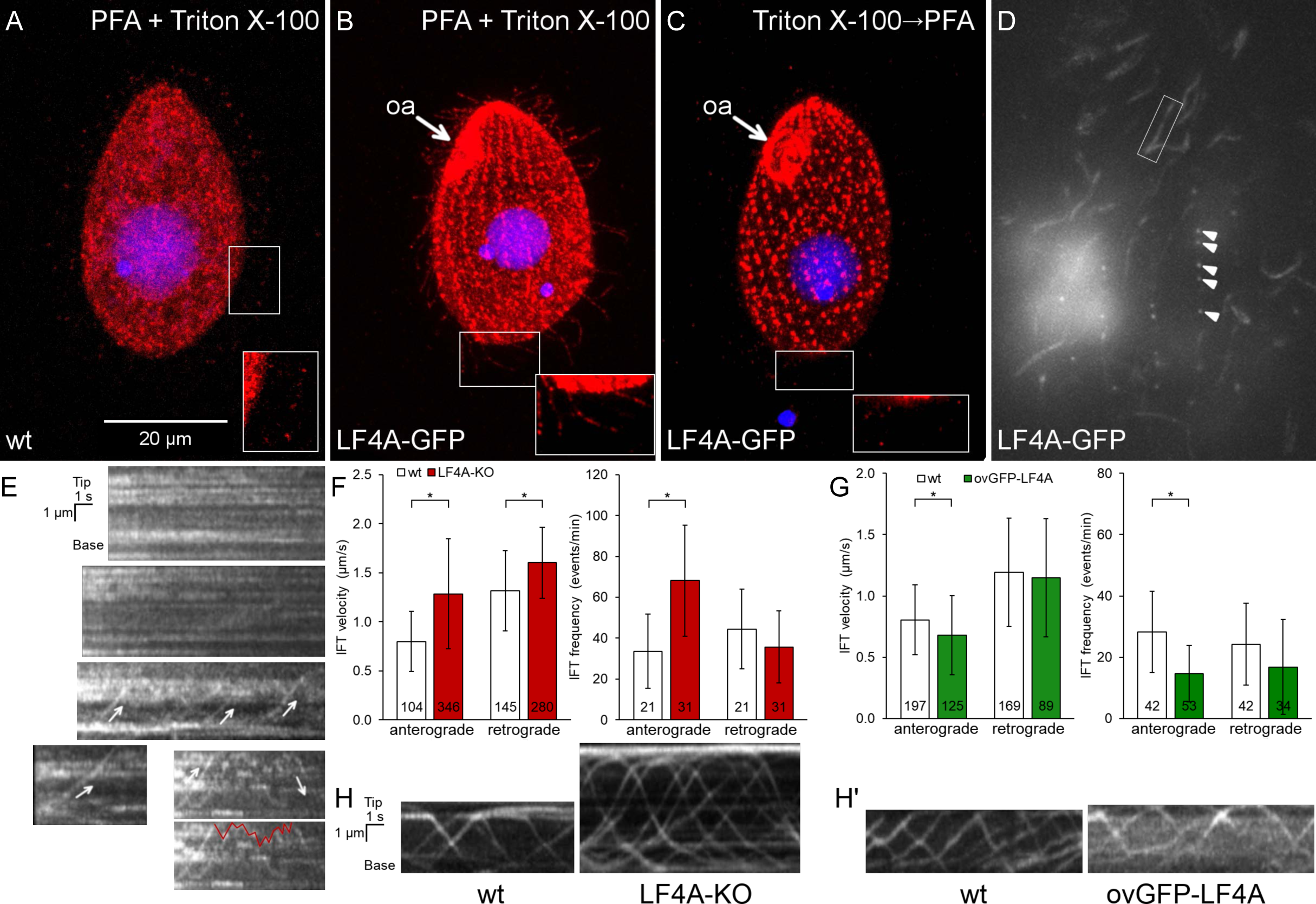
LF4A-GFP localizes to the basal bodies and along cilia, is transported by IFT and affects IFT. (A-B) A wild-type (negative control) cell (A) and a cell expressing LF4A-GFP under the native promoter (B) were subjected to simultaneous fixation/permeabilization (using a mixture of paraformaldehyde and Triton X-100) and stained with the anti-GFP antibodies (red) and DAPI (blue). In the negative control, the background red (anti-GFP) signal is in the cell body and occasionally at the tips of cilia (A inset). LF4A-GFP is present near the basal bodies and is weakly presents along the shafts of locomotory cilia (B inset). (C) An LF4A-GFP expressing cell that was first permeabilized (with Triton X-100) and then fixed (with paraformaldehyde) prior to immunofluorescence. Compared to (B), the LF4A-GFP signal remained strong near the basal bodies but is decreased in cilia. (D) A TIRFM image of a live LF4A-GFP cell. Arrowheads indicate several basal bodies within a locomotory row, the box highlights an example of a ciliary shaft. (E) Five representative example kymographs obtained from the TIRFM videos of LF4A-GFP cells. In most cilia, LF4A-GFP particles are stationary and some move with velocities similar to IFT velocities (arrows) or diffuse (an example of a diffusion event is marked with a red line on the duplicate of the bottom right kymograph). (F) The IFT velocities (left) and IFT event frequencies (right) in cilia of either wild-type or LF4A-KO cells that express DYF1-GFP. (G) The IFT velocities (left) and event frequencies (right) in cilia of either wild-type or GFP-LF4A-overexpressing cells. Both strains were exposed to Cd^2+^ for 3 hours. In (F) and (G), the sample sizes are indicated, asterisks mark significant differences (one way Anova p<0.01), error bars represent SDs. (H) Examples of kymographs of GFP-DYF1 in cilia of either wild-type or LF4A-KO cells corresponding to the data shown in panel F. (H’) Examples of kymographs of GFP-DYF1 in either wild-type or GFP-LF4A overproducing cells corresponding to the data shown in panel G. Abbreviations: oa, oral apparatus; PFA, paraformaldehyde.

The apparent paralog of LF4A, LF4B (Fig 1A) does not appear to be associated with cilia. LF4B-GFP was not detectable in the vegetatively-growing cells (S1A Fig). In the early phase of conjugation, LF4B-GFP localized to the junction between the two mating cells (S1B-S1C Fig), in a pattern that did not correspond to the positions of cilia in the vicinity of the conjugal junction [69-71]. Consistently, while the mRNA of *LF4A* is abundant in the vegetatively-growing cells, the mRNA of *LF4B* is present above the background only during the early stage of conjugation (S1D Fig, [72]). Likely, *Tetrahymena* has only a single cilia-associated LF4/MOK, LF4A.

### LF4A kinase activity shortens cilia and downregulates IFT

We overproduced GFP-LF4A, using the Cd^2+^-inducible MTT1 promoter [73]. After 6 or more hours of exposure to added Cd^2+^, GFP-LF4A strongly accumulated at the bases of both oral and locomotory cilia, all cilia shortened to became stumps (Fig. 3B compare to the GFP control in Fig 3A, Fig 3F), and the cells became paralyzed. In addition, based on TIRFM, overproduced GFP-LF4A decorated two non-ciliary microtubule-based structures: longitudinal microtubule bundles and contractile vacuole pores (S2A Fig, left panel), which suggests that LF4A has a microtubule-binding affinity. With time, the GFP-LF4A-overproducing cells became excessively large and misshaped (Fig 3D), indicating defects in cytokinesis. This is not surprising because in *Tetrahymena* locomotory cilia are required for the scission of daughter cells at the end of cytokinesis [74].

**Figure 3.**
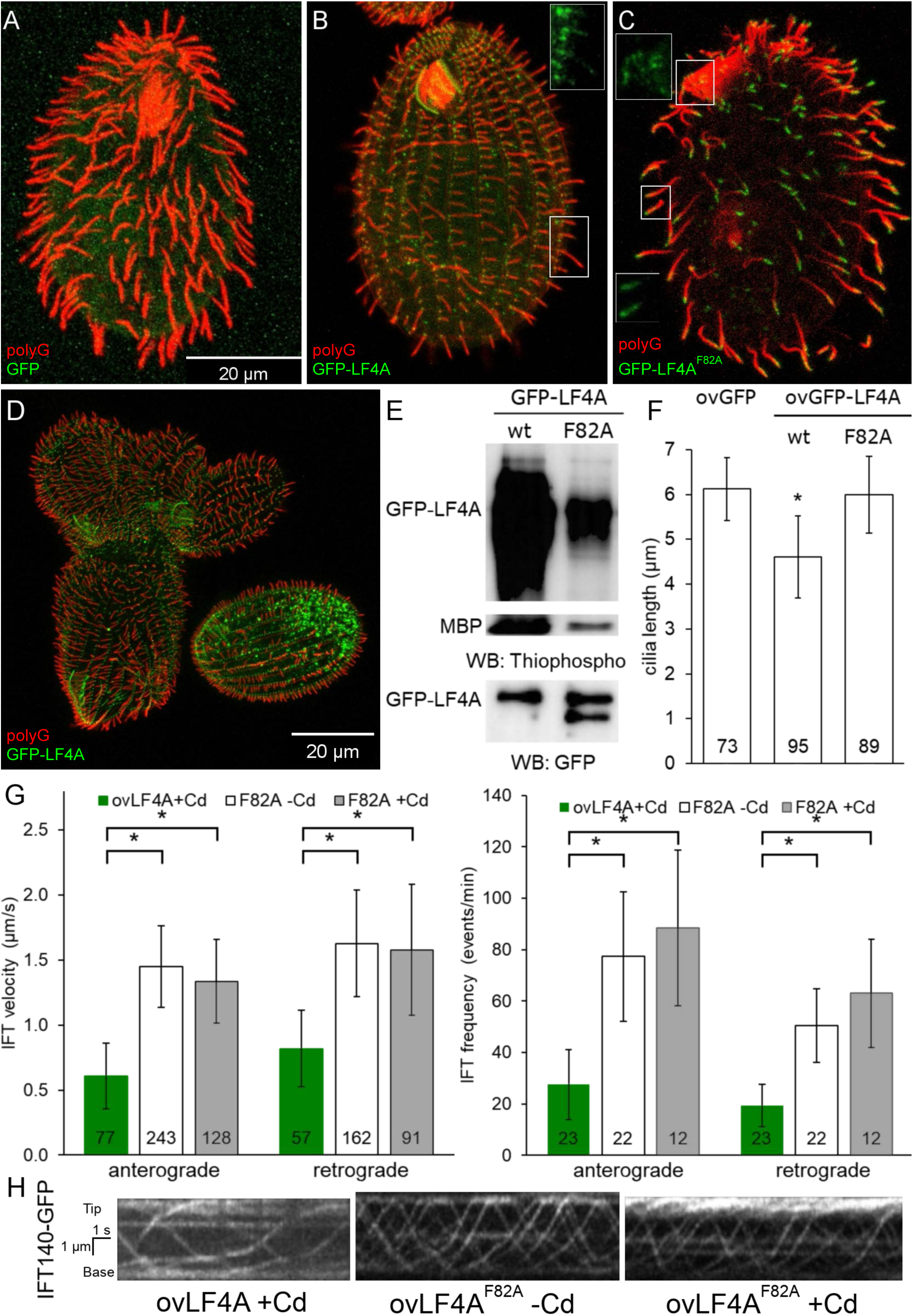
Excessive LF4A kinase activity shortens cilia and decreases IFT. (A-C) Cells (treated with 2.5 µg/ml CdCl_2_ for 6 hours) overexpressing either GFP (A), GFP-LF4A (B) or GFP-LF4A^F82A^ (C) showing GFP fluorescence (green) and labeled by the anti-polyG tubulin antibodies (red). The shortening of cilia is evident only in the cell expressing GFP-LF4A (B) where it is enriched at the basal bodies. GFP-LF4A^F82A^ accumulates at the tips of cilia (C). The insets in B-C show the green signal of GFP-LF4A alone at higher magnification. (D) GFP-LF4A-overexpressing cells after overnight induction with Cd^2+^, have short cilia, are large and irregular in shape, consistent with cytokinesis defects. (E) The results of an *in vitro* kinase assay with either GFP-LF4A or GFP-LF4^F82A^. Each protein was overexpressed in *Tetrahymena* and immunoprecipitated on anti-GFP-beads. The beads were incubated with the recombinant myelin basic protein (MBP) and ATP-γ-S. The phosphorylated products were detected on a western blot probed with the anti-thiophosphorylation antibody 51-8; the upper and middle panels show sections of the same blot containing autophosphorylated GFP-LF4A and MBP, respectively. The same amounts of IP inputs were analyzed on a western blot probed the anti-GFP antibodies and shown in the bottom panel (WB: GFP); the lower band in the right lane likely is a proteolytic degradation product of GFP-LF4A. (F) The lengths of locomotory cilia of cells that overproduce (for 3 hours) either GFP, GFP-LF4A or the kinase-weak GFP-LF4A^F82A^. The cilia lengths in the GFP and GFP-LF4A^F82A^ cells were not significantly different (p > 0.01), while the GFP-LF4A cilia were significantly reduced (∼75% of the length of GFP controls, one way Anova test, p < 0.01). Sample sizes are indicated, error bars represent SDs. (G) The anterograde and retrograde IFT velocities in cilia of cells that express IFT140-GFP and overexpress either mCherry-LF4A or mCherry-LF4A^F82A^. The IFT speeds are significantly reduced in mCherry-LF4A as compared to mCherry-LF4A^F82A^ overexpressing cells (exposed to added Cd^2+^ for 3 hours). The sample sizes (numbers of tracks measured) are indicated, asterisks indicate statistically significant differences (one way Anova, p<0.01), error bars represent SDs. (H) Examples of kymographs of cilia in cells expressing IFT140-GFP in different genetic backgrounds and conditions corresponding to the data shown in panel G. Scale bar: 1 µm × 1 s.

*In vitro*, LF4 of *Chlamydomonas* phosphorylates a generic substrate of serine/threonine kinases, myelin basic protein (MBP) and autophosphorylates [39]. Likewise, GFP-LF4A pulled down from overproducing *Tetrahymena*, phosphorylated MBP and itself *in vitro* (Fig 3E). An overproduced GFP-LF4A with a substitution of the conserved F82 (gatekeeper residue), GFP-LF4A^F82A^, had greatly reduced kinase activity *in vitro* (Fig 3E) and did not shorten cilia *in vivo* (Fig 3C and Fig 3F). While the overproduced GFP-LF4A accumulated at the ciliary bases (Fig. 3B and S2A Fig left panel), the kinase-weak GFP-LF4A^F82A^ accumulated at the tips of cilia (Fig 3C and S2A Fig middle panel). Overproduced mCherry-LF4A^F82A^ accumulated in cilia, but unlike GFP-LF4A^F82A^ was not enriched at the ciliary tips (S2A Fig right panel, S2C Fig middle and right panel). Thus, the tip enrichment of GFP-LF4A^F82A^ could be an artifact of the epitope tag, possibly caused by oligomerization of GFP [75]. Inside cilia, the particles of overproduced mCherry-LF4A and mCherry-LF4A^F82A^ were either immobile or moved with the IFT trains (S2B Fig). Thus, the kinase activity of LF4A is required for its cilia-shortening activity but is not required for its entry into cilia, anchorage or transport by IFT.

It is intriguing that only the kinase weak and not the active version of GFP-LF4A accumulates at the ciliary tips (Fig 3B-C, S2A Fig). The overproduced GFP-LF4A may fail to build up at the ciliary tips, if its kinase activity blocks the anterograde IFT of cargoes, including GFP-LF4A itself. We thus examined how the levels of LF4A affect IFT. In cells expressing a tagged IFT subcomplex B protein, GFP-DYF1/IFT70 [76, 77], the loss of LF4A significantly increased the velocities of both the anterograde and retrograde IFT (Fig 2F and 2H). Overexpression of GFP-LF4A mildly decelerated the anterograde but not retrograde IFT (Fig 2G and 2H’). Overproduced mCherry-LF4A (but not mCherry-LF4A^F82A^) decreased the velocities of both the anterograde and retrograde IFT based on imaging of a tagged IFT subcomplex A subunit IFT140 [19, 78], IFT140-GFP (Fig 3G left panel, Fig 3H). In addition, the loss of LF4A significantly increased the frequency of the anterograde (but not the retrograde) IFT events (Fig 2F right panel). Overexpression of GFP-LF4A strongly reduced the IFT event frequencies in the anterograde direction in cells with GFP-DYF1 reporter (Fig. 2G right panel, fig 2H’) while overexpression of mCherry-LF4A (but not mCherry-LF4A^F82A^) decreased IFT frequency in both directions in cells expressing IFT140-GFP (Fig 3G right panel, Fig 3H). The inhibitory impact of overproduced LF4A on IFT frequencies was likely underestimated, because many cilia in these cells lacked detectable IFT motility and very short cilia could not be analyzed (S2 and S3 movies). Overall these observations indicate that LF4A inhibits IFT by reducing both the frequency and velocity of IFT trains.

### Identification of an *LF4A* interactor, CDKR1

To find potential interactors of LF4/MOK, we performed a genetic screen for suppressors of GFP-LF4A overexpression, taking advantage of the resulting cell paralysis. We introduced a GFP-LF4A transgene operating under the MTT1 promoter and linked to the *neo5* marker (ovGFP-lf4a allele) into the (germline) micronucleus, by replacing the native *LF4A* locus (Fig 4A and S3 Fig). The transgene-carrying strain was mutagenized with nitrosoguanidine and subjected to self-fertilization by uniparental cytogamy [79]. This procedure generates whole genome homozygotes, each derived from a single, diploidized meiotic product of the parent cell, and thus allows for isolation of recessive and dominant mutations. Cd^2+^ was added to the mutagenized progeny to induce overexpression of GFP-LF4A and to paralyze the non-suppressed progeny. Suppressors were isolated based on their capacity to swim to the top of test tubes (Fig 4B). Five independent suppressor clones (designated as F0 generation clones) were isolated, while none were found among the progeny of a similar number (∼3 × 10^7^) of non-mutagenized cells. To distinguish between the extragenic and intragenic suppressions, we tested each suppressor mutation for linkage with the transgene-coupled *neo5* marker that confers resistance to paromomycin. The F0s were crossed to a wild type and the F1 heterozygous progeny were used to generate F2s by self-fertilization. Tight linkage (essentially 0% recombinants) was expected for an intragenic suppressor mutation while a completely unlinked single suppressor mutation would yield a 1:1 ratio of recombinant to parental F2 genotypes (Fig 4C and S3 Fig). Four suppressor clones (SUP2,3,4 and 5) were judged to be intragenic, and one suppressor clone (SUP1) was judged to be extragenic, based on the ∼1:2 ratio of the parental versus recombinant F2 phenotypes (the excess of recombinants could be spurious, due to unequal growth rates of suppressed and unsuppressed F2 progeny).

**Figure 4.**
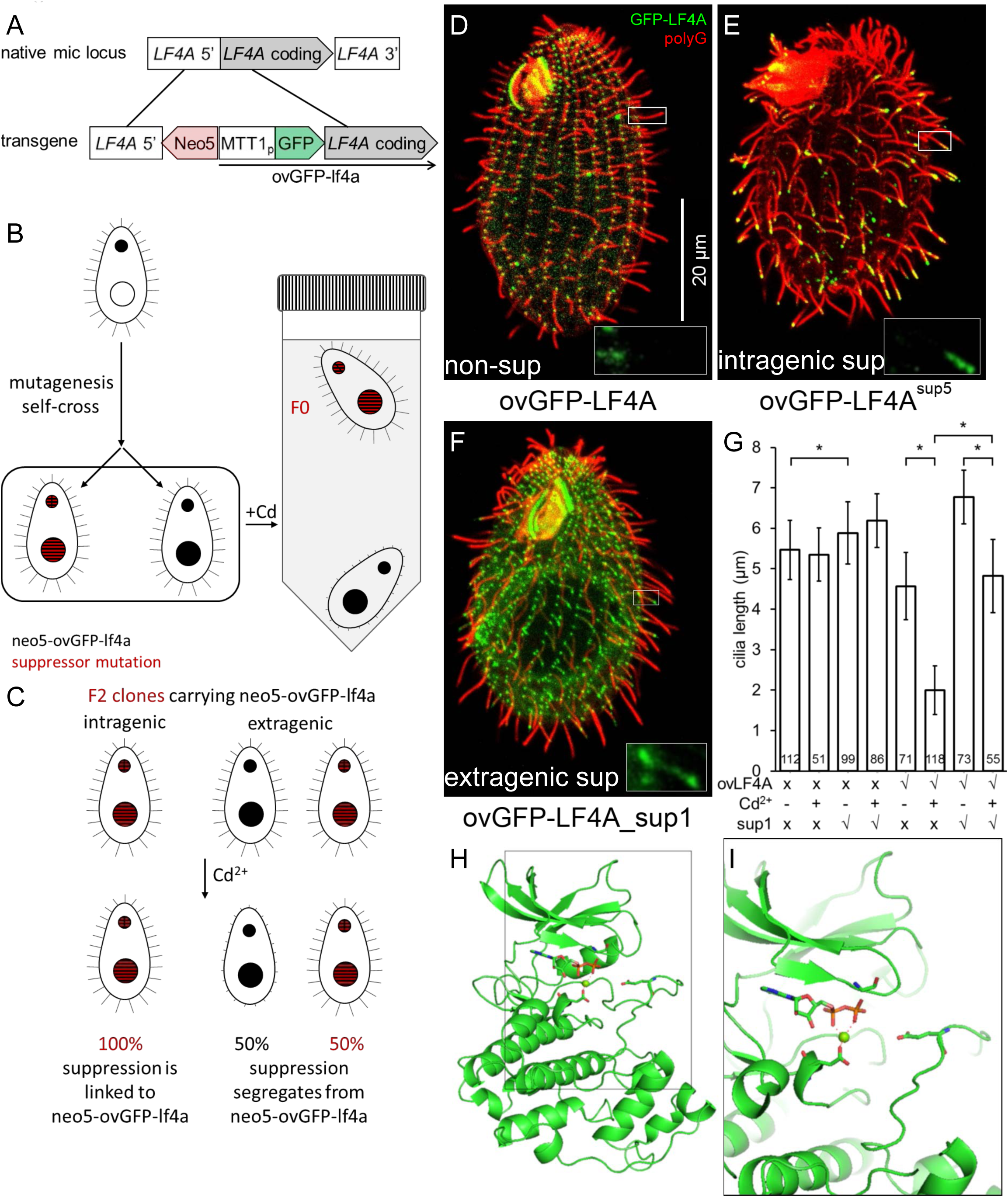
Isolation of intragenic and extragenic suppressors of GFP-LF4A overexpression. **(A-C)** The pipeline used for identification of intragenic and extragenic suppressors of overexpression of GFP-LF4A. (A) The structure of the transgene that was placed in the micronucleus. The transgene uses MTT1 promoter to express GFP-LF4A. A *neo5* cassette is closely linked. The transgene replaces the endogenous *LF4A*. (B) An outline of the procedure for generating suppressors. A heterokaryon with the ovGFP-lf4a transgene in the micronucleus (solid black) was subjected to mutagenesis and a self-fertilizing cross. The homozygous progeny was selected based on paromomycin resistance conferred by *neo5* and some progeny clones may carry a suppressor mutation (red stripes). The progeny cells were treated with Cd^2+^ in tubes kept in vertical position. The progeny clones that lack a suppressor mutation shorten cilia and sink to the tube bottom. The suppressors (F0) remain motile and accumulate near the top of the tube due to negative gravitaxis. (C) The principle of testing whether the suppression is intra- or extragenic (for details see S3 Fig). The F1 clones were subjected to a self-cross and the pm-r F2 progeny clones were isolated. An intragenic suppressor gives pm-r F2 progeny clones that are 100% motile (suppressed), as the suppression is linked to the transgene. An extragenic suppressor generates F2 pm-r clones that are either suppressed (motile) or not (paralyzed). (D-F) Cells that were exposed for 6 hours to Cd^2+^ to induce GFP-LF4A, subjected to immunofluorescence to reveal GFP (green) and polyG tubulin (red). Insets show the GFP-LF4A signal alone in examples of cilia at a higher magnification. (D) A non-mutagenized non-suppressed cell; GFP-LF4A is enriched at the bases of cilia. (E) An intragenic suppressor cell (SUP5); GFP-LF4A is enriched at the tips of both oral and locomotory cilia. (F) An extragenic suppressor cell (SUP1); GFP-LF4A is prominent at the bases, at the distal ends and along the ciliary shafts in short (presumably assembling) cilia. (G) The locomotory cilia length of F2 clones of four genotypes without and with 6 hours Cd^2+^ treatment. Sample sizes (number of cilia measured) are indicated, asterisks indicate a statistically significant difference (one way Anova p<0.01), error bars indicate SDs. (H) A 3D predicted structure of the kinase domain of LF4A based on homology-directed modeling using CDK of *Cryptosporidium* (Chain A of PDB 3NIZ) as template. (I) A zoomed-in view of a region of the structure showing three locations of substitutions (shown as sticks) found in the intragenic suppressors sup3 (E132K), sup4 (G13S) and sup5 (E160K).

When the four intragenic suppressors were exposed to Cd^2+^ to induce overproduction of GFP-LF4A, SUP2 lacked a GFP signal while SUP3, SUP4, SUP5 had a strong GFP-LF4A signal at the tips of cilia (Fig 4E compare to the non-suppressed cell in Fig 4D, S4 Fig), as seen earlier for GFP-LF4A^F82A^ (Fig 3C). The extragenic suppressor, SUP1, had a GFP-LF4A signal at the ciliary bases, but also the length and at the tips of short (presumably assembling) cilia (Fig 4F).

Sanger DNA sequencing of the ovGFP-LF4A transgene in SUP2 revealed multiple mutations in the MTT1 and GFP portions of the transgene, consistent with the lack of GFP fluorescence. SUP3, SUP4 and SUP5 carried single point mutations, predicted to result in E132K, G13S and E160K substitutions, respectively, in the kinase domain of LF4A. A homology-based model of the LF4A kinase domain (using the 3D structure of CDK of *Cryptosporidium* (Chain A of PDB 3NIZ) as a template [80, 81]) revealed that all three affected amino acids are adjacent to the kinase active site (Fig 4H, 4I). While it is not clear how these mutations affect LF4A, in other kinase types substitutions at the positions equivalent to G13 and E132 are associated with diseases [82].

To identify the causal mutation in the single extragenic suppressor SUP1, we used comparative whole-genome sequencing as recently described [83]. A number of independent (meiotic segregant) F2 clones, all derived from a single sup1/SUP1^+^ heterozygote, were combined into a suppressed and a non-suppressed pool (Fig 5A and 5B) and the pooled genomic DNAs were sequenced. The sequence variants found in the suppressed pool were subjected to bioinformatic subtractions (to remove variants also found in the unsuppressed pool and in other unrelated strains) and filtering (for nitrosoguanidine-type mutations [84]) (Fig 5C). These steps yielded three variants, each located on a different micronuclear chromosome (S1 Table). We alsoo used the “allelic composition contrast analysis (ACCA)” to plot the frequency of variant cosegregation with the suppression phenotype along each of the five micronuclear chromosomes [83]. A single peak of linkage was present on the micronuclear chromosome 3 at a bp location between 9 to 10 Mb (Fig 5E), a region that intersected with one of the three variants identified by subtractions and filtration: the T to C mutation on macronuclear scaffold 8254401 at bp location 105680, in the gene *TTHERM_01080590*, which encodes a kinase. The mutation changes the predicted stop into a tryptophan codon and ads a “WIRNLLILNG” sequence to the otherwise normal C-terminus of TTHERM_01080590 protein (Fig 5D). Based on a kinase profiling search [85, 86], TTHERM_01080590 is a CDK-related kinase and therefore we named the *TTHERM_01080590* gene *CDKR1* (<u>c</u>yclin-<u>d</u>ependent <u>k</u>inase-<u>r</u>elated 1). Among several metazoan species analyzed, CDKR1 is most similar to DYF-18 (Fig 1A), a known cilia-associated CDK-related kinase of *C. elegans*, which was proposed to be a homolog of LF2/CCRK CDK kinases [46, 48]. While our phylogenetic analysis does not support either DYF-18 or CDKR1 as orthologs of LF2/CCRK, we note that DYF-18, CDKR1, LF2 and CCRK are all CDK-type kinases that lack the cyclin-binding motif, PSTAIRE, characteristic of the canonical CDKs that regulate the cell cycle (S5C Fig), and are all associated with cilia where they act upstream of RCKs (see below and [40, 43, 47, 48, 64]).

**Figure 5.**
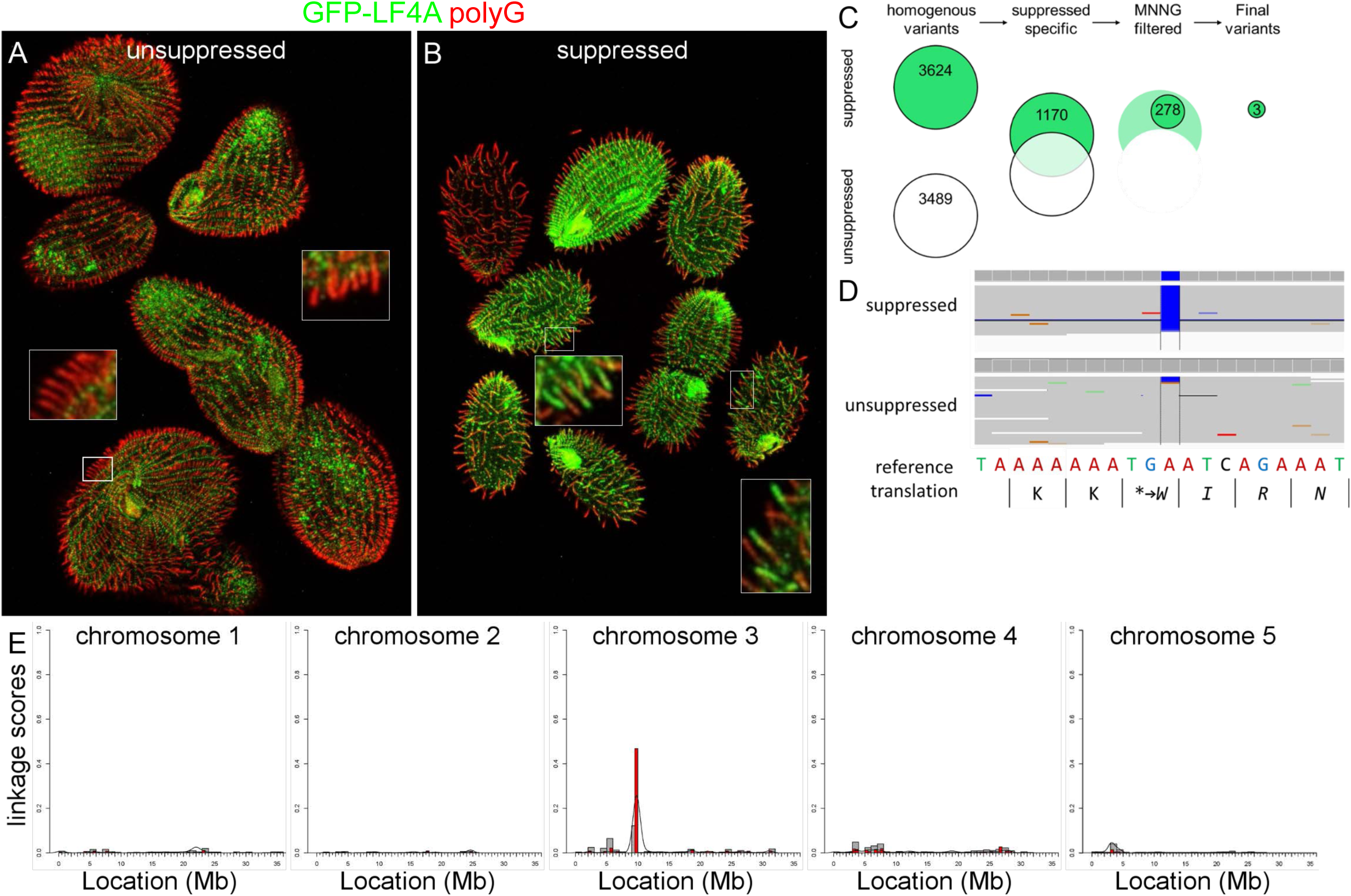
The extragenic suppressor clone SUP1 has a mutation in *CDKR1*. (A-B) Immunofluorescence images of (unsuppressed and suppressed) pools of F2 progeny derived from a single sup1/SUP1^+^ (after an overnight exposure to Cd^2+^) that were subsequently subjected to whole genome sequencing. Note the patterns of GFP-LF4A inside cilia: the base accumulation in the non-suppressed pool and the ciliary shaft and tip signals in the suppressed pool (see insets for higher magnifications). (C) The results of variant subtraction and filtering based on the alignment of sequencing reads to the macronuclear reference genome. Among the 278 variants consistent with nitrosoguanidine (MNNG) mutagenesis, three candidate variants affect a gene product and have a high fraction of reads supporting the alternative base in the mutant pool (see S1 Table). (D) An IGV browser view of the macronuclear genome sequence of *TTHERM_01080590* (*CDKR1*) that contains the variant scf_8254401:105680 T to C. This point mutation, supported by 100% of the sequencing reads from the mutant pool, changes the stop codon and adds a short peptide to the C-terminus of the predicted product. (E) An allelic composition contrast analysis of the variant co-segregation across all micronuclear chromosomes. The normalized linkage scores show the difference in the allelic composition between the mutant and the wild-type pool at each variant site. This reveals a cluster of variant co-segregation at 9-10 Mb on the micronuclear chromosome 3.

### CDKR1 is a negative regulator of cilium length that activates LF4A

In an otherwise wild-type background, the *cdkr1*^*sup1*^ allele mildly increased cilium length (Fig 4G). In cells overproducing GFP-LF4A, the *cdkr1*^*sup1*^ allele partially suppressed the shortening of cilia (Fig 4G). To clarify whether *cdkr1*^*sup1*^ is a gain or loss-of-function allele, we produced a strain with a null allele, CDKR1-KO. The locomotory cilia of CDKR1-KO cells were much longer than those of the wild-type or CDKR1^sup1^ cells (Fig 6C compare with 6A, 6E, 4G). Thus, likely the *cdkr1*^*sup1*^ allele is a hypomorph.

**Figure 6.**
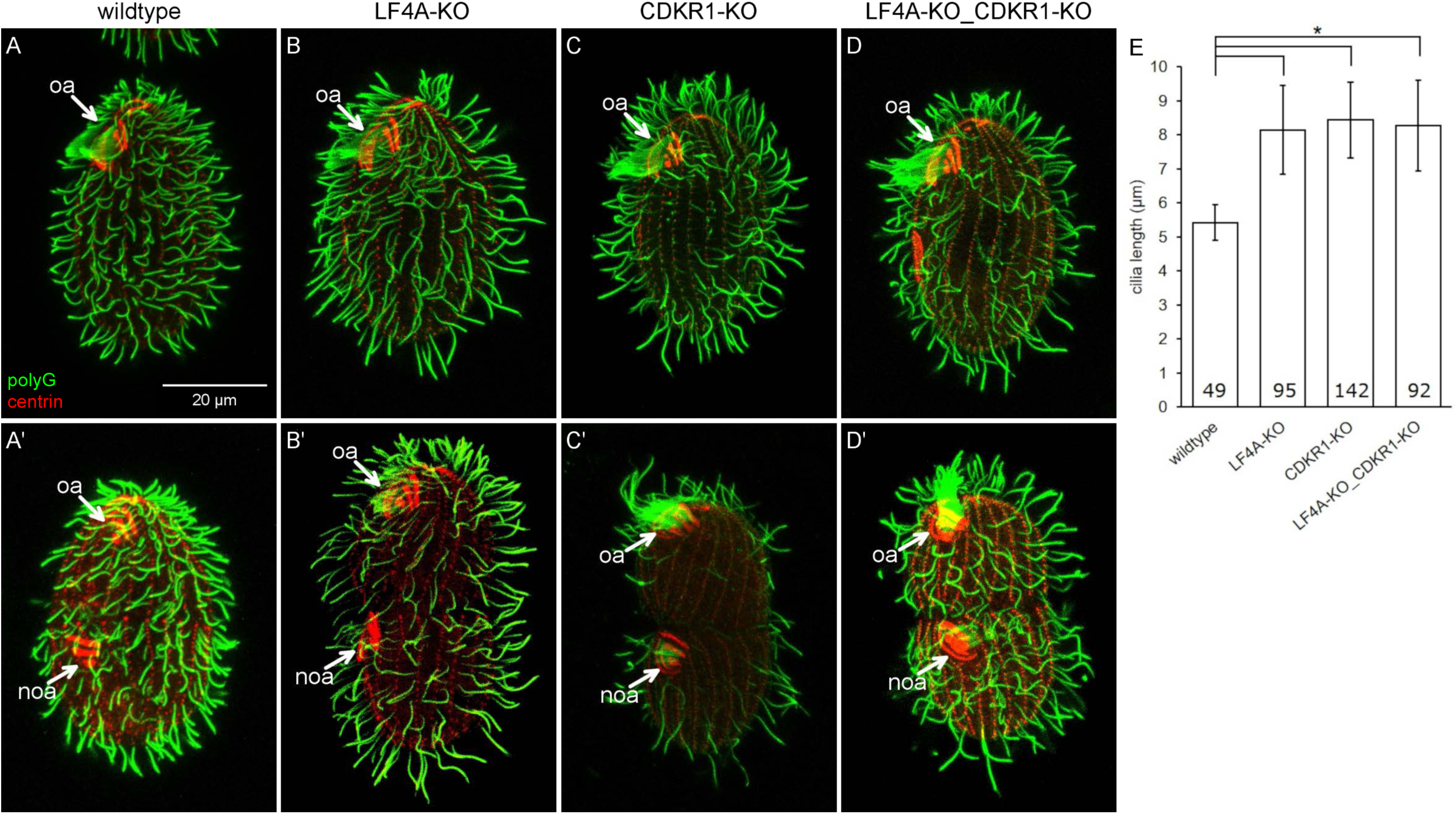
A loss of CDKR1 lengthens both the locomotory and oral cilia. (A-D’) Wildtype (A and A’), LF4A-KO (B and B’), CDKR1-KO (C and C’) and double knockout LF4A-KO_CDKR1-KO (D and D’) cells that are either in interphase (top panels) or dividing (bottom panels). The cells were stained with the anti-polyG (green) and anti-centrin antibodies (red). (E) The locomotory cilia lengths in different backgrounds. The cilia length in LF4A-KO, CDKR1-KO and the double knockout strain are similar (p > 0.01) and all are significantly longer than the wild-type cilia (one way Anova, p < 0.01). Sample sizes are indicated, error bars indicate SDs.

The CDKR1^sup1^ protein has 10 extra amino acids at the C-terminus but is otherwise normal, and thus the effect of the sup1 mutation was unclear. We used homology modeling to predict the structure of CDKR1 and its sup1 version. The closest 3D structure available is the human CDK2 (PDBID: 2IW8) [87], to which we could align most of CDKR1 (I12-N308). We attempted to model the remaining 23 (33 in sup1) C-terminal amino acids without a template. A Jpred [88] secondary structure prediction indicated that the LKKWIRNLL peptide in sup1 protein forms an alpha-helix. Thus, the WIRNLLILNG extension may enlarge the contact between the C-terminal tail of CDKR1 (that lies on the surface of the kinase domain) and the catalytically important C-helix (S5A Fig). As mentioned earlier, unlike the canonical CDKs that regulate the cell cycle, CDKR1 (and other cilia-associated CDK-related kinases DYF-18, LF2 and CCRK) lacks the cyclin-binding motif, PSTAIRE (S5C Fig). The WIRNLLILNG extension may pack to the region of the C-helix where cyclin typically binds in the canonical CDKs (S5B fig). Thus, the C-terminal tail extension in the sup1 version may affect the C-helix conformation in the critical regulatory region that is important for kinase activation.

To explore further how CDKR1 may interact with LF4A, we compared the phenotypes of the respective null mutants. The locomotory cilia of CDKR1-KO cells were similar in length to those of LF4A-KO, and also more sparsely present in the posterior cell region (Fig 6A-C and 6E). Strikingly, while the oral cilia seemed unaffected in LF4A-KO, they were exceptionally long in CDKR1-KO (Fig 6C compare to 6A and 6B, marked with “oa”). The excessively long oral cilia were most striking in the old (anterior) oral apparatus of the dividing cells (Fig 6C’ compare to 6A’ and 6B’, the old oral apparatus marked with “oa”) Unlike the wild-type and LF4A-KO cells, the CDKR1-KO cells could be maintained long-term only on the specialized medium MEPP that supports proliferation of mutants deficient in phagocytosis [89], indicating that the oral cilia in CDKR1-KO were functionally compromised. Overall the phenotype of CDKR1-KO was more severe as compared to LF4A-KO. The two null alleles similarly affected the locomotory cilia but only the loss of CDKR1 lengthened the oral cilia. The double knockout (LF4A-KO_CDKR1-KO) cells had the phenotype similar to the single knockout CDKR1-KO, including long oral cilia (Fig. 6D-D’ and Fig 6E). To summarize, both LF4A and CDKR1 regulate the length of locomotory cilia (likely by acting in the same linear pathway, see below), while only CDKR1 significantly contributes to the length of oral cilia.

Next, we examined how the phenotype of overexpression of GFP-LF4A (shortening of cilia) is affected by a complete loss of CDKR1. We compared two strains with the ovGFP-LF4A transgene that were either otherwise wild-type (CDKR1^+^) or CDKR1-KO. Without Cd^2+^ treatment, the ovGFP-LF4A_CDKR1^+^ cells had normal length cilia, while the ovGFP-LF4A_CDKR1-KO cells had fewer and excessively long locomotory and oral cilia, as expected (Fig 7A and 7B). When GFP-LF4A overexpression was induced with Cd^2+^, cilia shortened in both strains, but to a different degree. While ovGFP-LF4A_CDKR1^+^ cells experienced a strong shortening of all cilia (locomotory and oral) (Fig 7C), in the ovGFP-LF4A_CDKR1-KO cells, both the locomotory and oral cilia shortened only partially and consequently had about a wild-type length (Fig 7D compare to 1B). Next, we tested whether overproduction of GFP-LF4A in the CDKR1-KO background normalizes the functionality of cilia, by examining the cell multiplication rate, that in *Tetrahymena* is dependent on the health of both oral and locomotory cilia [65]. As expected, without added Cd^2+^, the ovGFP-LF4A_CDKR1-KO cells grew more slowly than the ovGFP-LF4A_CDKR1^+^ cells (Fig 7G). Remarkably, after addition of Cd^2+^, the multiplication rate pattern had inverted; the ovGFP-LF4A_CDKR1^+^ cells ceased to multiply, while the ovGFP-LF4A_CDKR1-KO cells multiplied faster (Fig 7G). Thus, a complete loss of CDKR1 is rescued by overexpression of LF4A. These observations argue that the major if not only function of CDKR1 is to positively regulate a kind of activity provided by LF4A.

**Figure 7.**
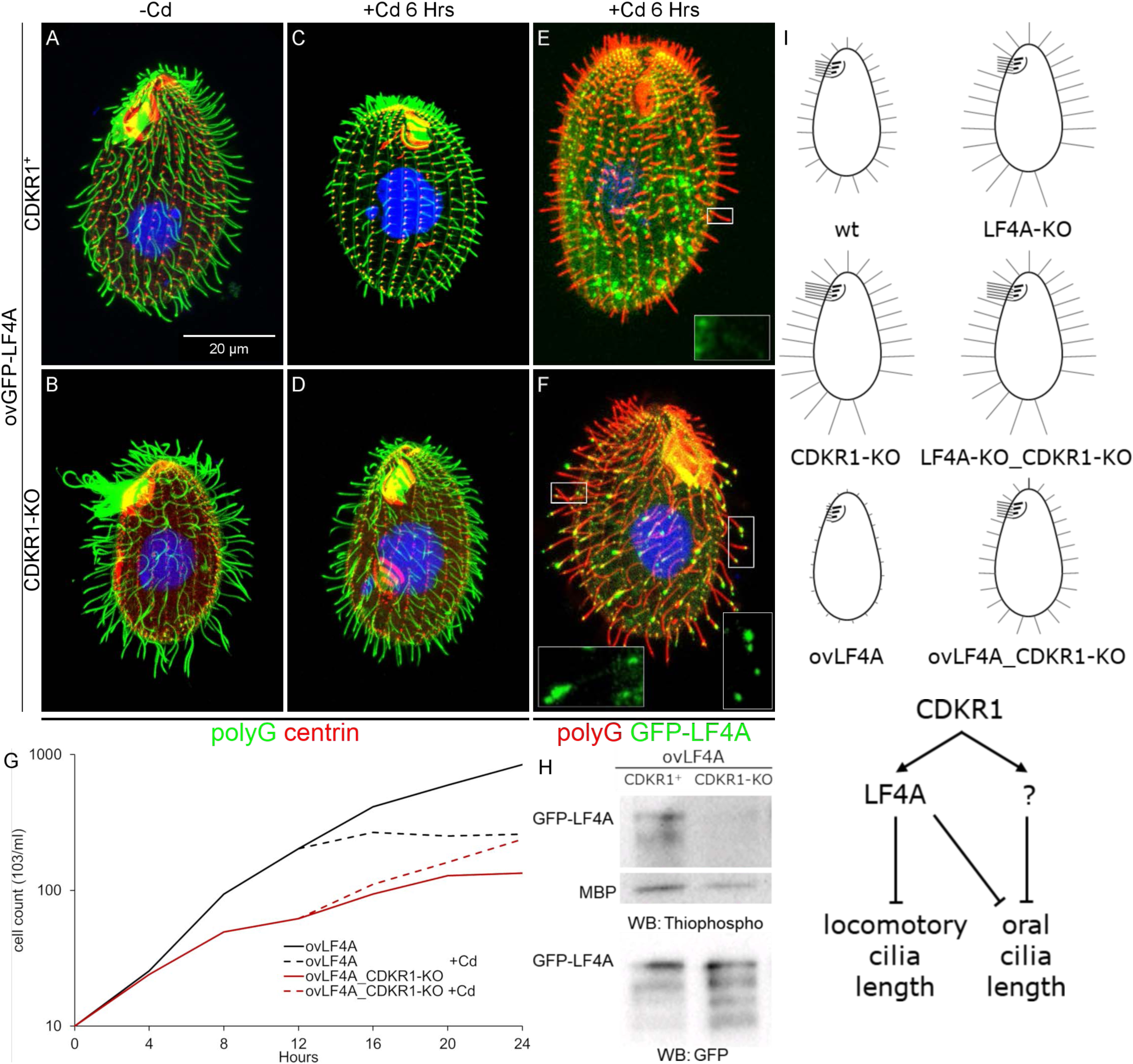
A complete loss of CDKR1 rescues the cilia shortening induced by GFP-LF4A overexpression and reduces the LF4A kinase activity. GFP-overproducing cells that have either wild-type CDKR1 (A, C, E) or are CDKR1-KO (B, D, F), imaged before (A and B) or after a 6 hours exposure to Cd^2+^ (C, D, E and F). In A-D, the cells were stained with the anti-polyG antibodies (green) and anti-centrin antibodies (red). In (E) and (F), the cells show a GFP-LF4A signal (green) and were stained with anti-polyG antibodies (red) antibodies. In cells lacking CDKR1, overproduced GFP-LF4A accumulated at the distal ends of cilia (F), indicating a reduced LF4A kinase activity. (G) The growth rates of multiple strains that overproduce GFP-LF4A and are either wild-type or CDKR1-KO. The cells were inoculated in SPPA media without Cd^2+^ (each data point averages 6 cultures), and an overexpression of GFP-LF4A was induced at 12 hours (each data point averages 3 cultures hereafter). (H) An *in vitro* kinase activity of overproduced GFP-LF4A isolated from two cells that are either otherwise (CDKR1^+^) and CDKR1-KO. The phosphorylated products were detected on a western blot probed with the anti-thiophosphorylation antibody 51-8 (the upper and middle panels show areas of the same blot containing the autophosphorylated GFP-LF4A and MBP, respectively). The same amounts of IP inputs were analyzed on a western blot probed the anti-GFP antibodies shown in the bottom panel (WB: GFP) the multiple bands likely are likely proteolytic degradation products of GFP-LF4A. (I) Top: a graphical summary of the phenotypes and genotypes. Bottom: a scheme of the likely pathway involving CDKR1, LF4A and another kinase, likely an RCK that acts downstream of CDKR1 in oral cilia.

One way how a loss of CDKR1 may affect the outcome of overexpression of GFP-LF4A is by decreasing its stability. Among three ovGFP-LF4A_CDKR1-KO clones (all derived from the same F1) that were phenotypically similar, the levels of overexpressed GFP-LF4A were highly variable (S6A Fig). Similarly, the levels of overproduced GFP-LF4A varied among several clones that all carried the *cdkr1*^*sup1*^ allele (S6B Fig). While the sources of this variability are unclear, it appears that the suppression phenotype does not strictly correlate with the levels of overproduced GFP-LF4A. Strikingly, GFP-LF4A overproduced in the CDKR1-KO cells, strongly accumulated near the tips of cilia (Fig 7F compare to Fig 7E), which is a phenocopy of the kinase-weak GFP-LF4A^F82A^ (Fig 3C and S4C-E). Also, GFP-LF4 was enriched at the tips of some cilia in the presence of the hypomorphic allele *cdkr1*^*sup1*^ but not in the wild-type background (Fig 4F compare to 4D). These observations suggest that deficiencies of CDKR1 reduce the kinase activity of overproduced GFP-LF4A. Indeed, overproduced GFP-LF4A pulled down from either the CDKR1-KO or cdkr1^sup1^ cells had reduced kinase activity *in vitro* as compared to the same protein from a wild-type background (Fig 7H and S6C). Thus, CDKR1 increases the kinase activity of LF4A. However, even without CDKR1, overproduced LF4A has residual kinase activity *in vitro* (Fig. 7H) and *in vivo* as it partially shortens cilia (Fig. 7D compare to 7B).

Under the native conditions, CDKR1 regulates the length of locomotory cilia, most likely by activating LF4A (based on the similarity of the null phenotypes in regard to the locomotory cilia, Fig 6). To regulate the length of oral cilia, CDKR1 may activate LF4A and one or more of the unstudied RCKs (Fig. 1A) that could be partially redundant with LF4A. That LF4A functions in both locomotory and oral cilia (despite a lack of effect of its loss on the length of oral cilia) is indicated by 1) the presence of LF4A-GFP near the basal bodies (Fig 2B, 2C), the cilia-shortening activity of GFP-LF4A (Fig 3B) and the enrichment of multiple kinase-weak variants of GFP-LF4A at the tips (Fig 3C and S4C-E) of both locomotory and oral cilia. To summarize, CDKR1 increases the kinase activity of LF4A to promote shortening of locomotory cilia and may act through LF4A and another RCK to shorten the oral cilia (Fig. 7I).

## Discussion

### Ciliary and non-ciliary roles of RCKs

LF4/MOK and DYF-5/MAK/ICK are two conserved subgroups of RCK kinases. While most ciliated lineages, including mammals, have both LF4/MOK and DYF-5/MAK/ICK, *C. elegans, Drosophila* and zebrafish lack LF4/MOK (Fig 1A); thus DYF-5/MAK/ICK can be sufficient for ciliary functions suggesting that the two subtypes of RCKs have closely-related activities. This is not the case of *Chlamydomonas* [39, 64], and *Tetrahymena* (this study) where LF4/MOK loss-of-function mutants have abnormally long cilia despite the presence of DYF-5/MAK/ICK homologs. To our knowledge the significance of the mammalian ortholog MOK has not been established yet. While most RCKs are linked to cilia, some non-ciliated species, including *Dictyostelium discoideum* and fungi, have RCKs (Fig 1A). In the budding yeast its RCK, Ime2, functions in meiosis and sporulation (reviewed in [90]). We show that among the two LF4/MOK kinases of *Tetrahymena*, LF4A regulates cilia, while LF4B is expressed during conjugation. Possibly, following the whole genome duplication [91], LF4A retained the ancestral ciliary function [92], while LF4B underwent neofunctionalization.

### RCKs and their CDK activators affect both cilium length and number

The *Tetrahymena* cells lacking either LF4A or its activator, CDKR1 (see below), have longer but also fewer locomotory cilia. In the mouse, a loss of ICK or its activator CCRK, in different cell types leads to either longer or shorter cilia [5, 42, 93]. Importantly, in *Chlamydomonas,* hypomorphic LF2 alleles confer longer cilia, but a null allele produces variable length (including shorter) cilia and inability to regenerate cilia after deciliation [47, 64, 94]. Loss of either RCKs or LF2/CCRK cause excessive accumulation of IFT materials in cilia, which could create roadblocks that reduce the efficiency of IFT [40-42, 47, 93, 95]. In the multiciliated cells, such as *Tetrahymena*, the excessively long cilia may deplete factors whose concentration is rate-limiting for ciliogenesis in other parts of the same cell. To summarize, the long, short or even absent cilia could be parts of a single phenotypic spectrum caused by an underlying defect of excessive cilia assembly.

### RCKs inhibit IFT

The key question is how LF4/MOK (and other RCKs) control cilium length. We show that overexpression of LF4A shortens cilia and decreases the IFT speeds and frequencies. These observations, together with the work by others (see below) indicate that LF4/MOK (and more broadly RCKs) act on cilium length by inhibiting IFT. In *Tetrahymena*, the anterograde IFT is needed for outgrowth of cilia from the basal bodies [96-98], while the retrograde IFT is not required for ciliogenesis (without retrograde IFT, *Tetrahymena* cells assemble cilia that are more variable in length but have a nearly normal average length [99]). Thus, the phenotype of overexpression of GFP-LF4A is a phenocopy of a loss of the anterograde but not the retrograde IFT, suggesting that LF4/MOK inhibits the anterograde IFT.

We show here that the LF4A activity reduces the velocities of IFT trains. In particular, the anterograde IFT direction was consistently affected by manipulations of LF4A in all assays used. In *Chlamydomonas* and mammalian cells, deficiencies in LF4/MOK did not change the IFT train velocities [38, 40, 100]. On the other hand, in mammalian cells, a depletion of ICK increased the rate of anterograde IFT and its overexpression reduced the rate of retrograde IFT, respectively [38]. However, in *C. elegans,* loss-of-function mutations of DYF-5 reduce IFT speeds in both directions [41, 46]. Thus, the effects of RCKs on IFT velocities are inconsistent across different models. In *Chlamydomonas,* the anterograde IFT velocity increases as the growing cilium lengthens [34]. It is therefore possible that at least some changes in the IFT velocities caused by manipulations of RCKs in different models are secondary to changes in cilium length.

RCKs could be controlling cilium length by reducing the frequency of IFT trains, which would reduce the delivery of precursors needed for assembly of cilia. In support of this model, we show that a loss LF4A increases the frequency of anterograde and retrograde IFT trains. We also show that overexpression of LF4A reduces the IFT frequencies. One caveat is that our imaging approach may not detect smaller IFT particles and it is already known that size of IFT particles changes (decreases) as the cilium grows [34]. In *Chlamydomonas* a loss of LF4 increases the amount of IFT motors entering cilia [35]. Thus our observations and those made in *Chlamydomonas* [35] indicate that LF4/MOK reduces the pool of IFT trains entering cilia. It is also well documented that cilia deficient in RCKs have elevated levels IFT proteins, including cilia of *Chlamydomonas* lacking LF4 [40] and cilia of mammalian cells [42, 95] and *C. elegans* [41] deficient in DYF-5/ICK/MAK. In the absence of RCKs, excessive anterograde IFT may not be balanced by the retrograde IFT, leading to accumulation of IFT proteins in cilia.

Another parameter that may contribute to the assembly rate is the cargo occupancy on IFT trains, which is known to be higher in growing cilia as compared to the steady-state or disassembling cilia [16, 17, 36]. While to our knowledge, the effects of RCKs on the IFT cargo occupancy have not been studied, in *Chlamydomonas,* a loss of LF2, a CDK kinase that acts upstream of LF4 [39, 40, 47, 64], increases the frequency of IFT trains carrying tubulin [16]. Here we also show that LF4A is both a regulator and a cargo of IFT. An overexpressed kinase-weak variant of GFP-LF4A accumulated at the ciliary tips while an active GFP-LF4A accumulated at the ciliary base, which agrees with the model that the kinase activity of LF4A inhibits the IFT-mediated transport of cargoes (including itself) to the tips of cilia. Similar observations were reported for ICK in mammalian cells [5, 38]. All these observations taken together are consistent with RCKs inhibiting transport of cargoes to the ciliary tip through inhibition of the anterograde IFT.

Several recent studies have linked cilium length kinases, including RCKs, to regulation of kinesin-2, the anterograde IFT motor. In *C. elegans*, loss-of-function mutations in DYF-5, and its likely upstream activator DYF-18, rescue the short cilia phenotype caused an autoinhibitory mutation in OSM-3 kinesin-2 subunit, indicating that DYF-5 and DYF-18 inhibit OSM-3 [46]. *In vitro*, ICK phosphorylates the tail of murine kinesin-2 motor subunit, KIF3A, at T674. While, in mammalian cells cilia are only mildly affected by T674A, mutating multiple phosphorylatable amino acids in the tail of KIF3A inhibits ciliogenesis in zebrafish [42]. In *Chlamydomonas*, CDPK1 phosphorylates the tail of kinesin-2 motor FLA8, on S663 *in vitro*. A phospho-mimicking mutant of FLA8 (S663D) lacks cilia, presumably due to inability of kinesin-2 to associate with IFT trains. While S663A mutation or depletion of CDPK1 both result in shorter cilia, CDPK1 also promotes the turnaround of IFT materials at the ciliary tip [60]. To summarize, the cilium-shortening influences of RCKs and CDPK1 may be mediated by phosphorylation and inhibition of kinesin-2 and the resulting reduction of anterograde IFT.

To our knowledge, we are first to image an RCK in live cells under near-native conditions. Strikingly, in the cilium, most of LF4A-GFP particles are stationary and scattered along the cilium length. A small subset of LF4A-GFP particles occasionally undergoes diffusion or moves with the IFT speeds as reported for overexpressed MOK and ICK in mammalian cells [38] and DYF-5 in *C. elegans* [41, 46]. LF4A may have a microtubule-binding ability, based on our observation that overproduced GFP-LF4A decorates non-ciliary microtubules. While our study suggest that ciliary LF4 is weakly anchored, in *Chlamydomonas* most of the ciliary LF4 remains associated with the axoneme after detergent extraction [40]. An anchorage to the axoneme could concentrate LF4/MOK near the passing IFT trains. The total exposure of IFT components to LF4/MOK could therefore be higher in a longer cilium. In this manner, an increased axoneme length may be translated into a proportionally decreased IFT activity, which would balance the rate of assembly with the rate of disassembly at steady state. Others have shown that the length of microtubules influences their properties by increasing the total amount of regulators landing on the polymer surface. For example, the end depolymerization rate is higher in a longer microtubule due to increased landing of depolymerizers that build up to a higher concentration at microtubule ends [101, 102]. As argued above, the activity of LF4/MOK decreases the IFT train entry rate. Because IFT trains enter at the ciliary base, a component of IFT trains would need to be reused in the subsequent IFT round, to convey a length-dependent feedback of the axoneme-anchored LF4/MOK. While many of the IFT components are replaced by fresh IFT proteins that arrive from the cell body, some (including IFT54) are partially recycled [103].

### CDK-related kinases (LF2/CCRK and DYF-18/CDKR1) are activators of RCKs

The first evidence that CDK-related LF2/CCRK kinases acts upstream of RCKs was an observation in *Chlamydomonas,* that an LF2 mutation is epistatic to an LF4 mutation [39]. The mammalian ortholog of LF2, CCRK, phosphorylates ICK and MAK on threonine of the <u>T</u>xY motif in the kinase activation loop [104-107]. In glioblastoma cells, overexpression of ICK inhibits ciliogenesis and this effect is suppressed by either a CCRK knockdown or by mutating T157 in the TxY motif of ICK [43]. A recent study found that in *Chlamydomonas* LF4 is phosphorylated at T159 of the TxY motif, and this phosphorylation requires LF2, suggesting that LF2 phosphorylates T159 of LF4 [40]. Here we used an unbiased approach to identify a CDK-related protein, CDKR1, as an activator of LF4A. CDKR1 resembles the LF2/CCRK and DYF-18. All these cilia-length regulating kinases are structurally similar to the canonical CDKs but lack the PSTAIRE cyclin-binding motif (S5C Fig). By analogy to other cilia-associated CDK kinases, CDKR1 may activate LF4A by phosphorylating the threonine in the TxY motif. Fu and colleagues showed that ICK is also weakly activated by autophosphorylation of tyrosine in the TxY motif [105]. Consistently, we show that LF4A is weakly active even without CDKR1 *in vitro* and *in vivo*. That overproduced LF4A rescues the cilia defects caused by a loss of CDKR1 indicates that the major function of CDKR1 is to activate LF4A or a similar activity provided by another kinase. Likely, CDKR1 acts by activating LF4A in the locomotory cilia and as discussed above, in addition to LF4A, activates another RCK in oral cilia. Our observations suggest that multiple RCKs are differentially utilized in different types of cilia and perhaps also among cilia located at different positions. Future studies on the seven uncharacterized DYF-5/ICK/MAK kinases of *Tetrahymena* could provide further insights into how subsets of cilia are managed by multiple length-regulating kinases in a single cell.

## Materials and methods

### Phylogenetic analysis of cilium length kinases

The cilium length-associated and CDK members of the CMGC kinase group were identified in multiple species by reciprocal BLASTp searches using sequences of well-studied proteins. Sequences were aligned with ClustalX 1.82 [108] and corrected manually in SEAVIEW [109]. A neighbor-joining tree was calculated with the Phylip package (using SEQBOOT, PROTDIST, NEIGHBOR and CONSENSE) [110]. The tree was visualized using FIGTREE (http://tree.bio.ed.ac.uk/software/figtree/). The following are NCBI accession numbers, names and abbreviated names used for the alignments and the phylogenetic tree: XP_001030514.2 (TTHERM_01080590, CDKR1), NP_503323.2 (DYF-18_Ce), NP_002737.2 (MAPK3_Hs), XP_001018592.1 (TTHERM_00286770), EAR97232.2 (TTHERM_00483640), XP_001020875.2 (TTHERM_00411810), NP_524420.1 (CDK2_Dm), XP_001027728.1 (TTHERM_01035490), XP_0010219111.2 (TTHERM_01207660), NP_001777.1 (CDK1_Hs), NP_009718.3 (CDC28_Sc), XP_001698637.1 (CDK_Cr), AAF55917.1 (CG6800_Dm), ABK34487.1 (LF2_Cr), NP_001034892.1 (CCRK_Hs), XP_009302023.1 (CDK20_Dr), NP_490952.2 (CDK7_Ce), NP_001790.1 (CDK7_Hs), NP_001256291.1 (CDKL5_Ce), NP_001124243.1 (CDKL5_Dr), NP_001310218.1 (CDKL5_Hs), XP_001008848.2 (TTHERM_00185770), AGC12987.1 (LF5_Cr), EAR90584.2 (TTHERM_00122330), EAR90792.3 (TTHERM_00141000), NP_055041.1 (MOK_Hs), AAO86687.1 (LF4_Cr), EAR83896.4 (TTHERM_00822360, LF4B), EAR87368.2 (TTHERM_00058800, LF4A), EAR93150.1 (TTHERM_00450990), EAR95676.2 (TTHERM_00267860), EAR89127.2 (TTHERM_00576780), EAR90889.2 (TTHERM_00144940), EAR82017.1 (TTHERM_01347900), XP_001697865.1 (MAPK7_Cr), NP_001129786.2 (Dyf-5_Ce), NP_001260307.1 (MAPK7_Dm), XP_009295564.1 (MAK_Dr), XP_011512721.1 (ICK_Hs), NP_005897.1 (MAK_Hs), XP_647537_Dd.

### Strains and cultures and cilia regeneration

SPP medium [111] with antibiotics, SPPA [112] was used to grow *Tetrahymena* with exception of strains with severe cilia defects, which were maintained in MEPP medium [89] with 2 µg/ml dextrose (MEPPD) [25]. CdCl_2_ (2.5 µg/ml) was added to induce MTT1-driven overproduction of proteins [73]. Deciliation and cilia regeneration were done as described in [113].

### Gene disruption, native locus tagging and overexpression

#### Native locus-based epitope tags

To tag *LF4A* at the native locus, the plasmid pKIN13AnativeGFP [25] was modified to make a derivative to target the 3’ end of the *LF4A* coding region. To this end, 1.3 kb and 0.7 kb fragments of *LF4A* were amplified using the primer pairs: 5’-AATACCGCGGACTTTCAACCAAACAAAACTCA-3’, 5’-TATTACGCGTTACTTATTAAAAACTGGCTTTTTACC-3’ and 5’-AATAATCGATAAACTACTTTATAGCTGTTTGTTTTTGA-3’, 5’-TTATGAGCTCGTGAGTCTAAACCTCCAGCAG-3’ and cloned on the sides of a fragment consisting of a GFP coding region, a *BTU1* transcription terminator and a *neo3* cassette in reverse orientation.

We constructed a *neo5* cassette that is similar to the one published in [114]. To tag LF4B at the native locus, two 1.1 kb fragments were amplified using the primer pairs: 5’-ATAAGGGCCCGCAGCAGATGATAGTGGAG-3’, 5’-TATTGAGCTCCATAGCATGGTACAGGAATCG-3’, 5’-ATAACCGCGGTAAGTCTTTTTCAATGTTTATGC-3’, 5’-TATTCTCGAGGAAAAAGGCTGGCAAGCG-3’ and cloned on the sides of a fragment consisting of a GFP coding region, a *BTU1* transcription terminator and *neo5* in reverse orientation.

To engineer a plasmid for tagging IFT140 in the native locus, a 0.5 kb terminal fragment of a the IFT140 coding region (TTHERM_00220810) was amplified using primers 5’-AATA ACGCGTGTATTGAGTAATTAGAAACTAAGCTCAA-3’ and 5’-AATTGGATCCTTCTGGGACATCTTCTTCAATG-3’ and used to replace the corresponding part of pFAP43-GFP-neo4 [115] plasmid using MluI and BamHI sites. Next, a 0.9 kb fragment of the 3’ UTR of *IFT140* was amplified using primers 5’-AAATCTGCAGCTTCATAGTAACTGACTACATTTAAAA-3’ and 5’-AATTCTCGAGACAAGCCATGCGAAAATG-3’ and cloned into pFAP43-GFP-neo4 using PstI and XhoI sites. Next *neo4* was replaced by the *pac* cassette that confers resistance to puromycin [116]. The resulted plasmid pIFT140-GFP-pac enables native expression of a IFT140 with a C-terminal GFP tag separated from IFT140 by a short linker (GSGGGSGTG).

#### Gene knockouts

For disruption of *LF4A*, two 1.2 kb genomic fragments of *LF4A* were amplified using the primer pairs: 5’-TATTGGGCCCTAATTTTATGTGATAGTCTTTATG-3’, 5’-TTATCCCGGGTGATTATCTCTAAATATTAATGTC-3’, and 5’-TTAACTGCAGCAGATATATATGGGATAATATTTA-3’, 5’-ATATCCGCGGTTTAGGAGTATATTTTCATAGTAT-3’ and cloned on the sides of *neo4* [117]. The germline-based total homozygotes were tested for the absence of the targeted *LF4A* fragment using a diagnostic PCR with primers: 5’-GTTTCGCCTCATCCTCACAT-3’ and 5’-AGAGAGATAATATGCAGGGCG-3’.

To disrupt *CDKR1*, the targeting homology arms were amplified with the following primer pairs: 5’-TATTGAGCTCAAATTTGAGGCACTACATTC-3’, 5’-TATTCCGCGGATTACCAAGCAAATCAG-3’ and 5’-TATTAAGCTTCATAAGCAAAAATAAAATGCC-3’, 5’-TATTATCGATGTAAAACTGAGAGCATTTGC-3’ and cloned on the sides of *neo5*. The loss of the targeted part of *CDKR1* was confirmed in homozygotes using diagnostic primers: 5’-TTTAAAGATGACTCTGTACC-3’ and 5’-CTGCAAGAGACTTGTATGC-3’ that amplify the targeted sequence.

#### Overexpression

For overexpression of GFP-LF4A at the *BTU1* locus, a 2.7 kb *LF4A* coding sequence was amplified using primers: 5’-TATTACGCGTCATGAACTAATATAAATTG-3’, 5’-TATTGGATCCTCATTACTTATTAAAAAC-3’ and cloned into pMTT1-GFP [118]. For overexpression of the kinase-weak GFP-LF4A^F82A^ variant, the pMTT1-GFP-LF4A plasmid was subjected to site-directed mutagenesis using the QuikChange Lightning kit (Agilent 210518) with the primers 5’-CAGGACGTTTGGCACTAGTGGCTGAATTGATGGATCAGAACC-3’ and 5’-GGTTCTGATCCATCAATTCAGCCACTAGTGCCAAACGTCCTG-3’.

For overexpression of GFP-LF4A at the native (*LF4A*) locus, a 1.3 kb 5’ UTR fragment of *LF4A* was amplified with primers: 5’-AATAGAGCTCATTAAGATCTCCTAACATGGAAT-3’, 5’-TATTCCGCGGCTTCTCTGAGTAGCTTCAAACAA-3’. Next, a 3.5 kb fragment of GFP-LF4A from pMTT1p-GFP-LF4A was amplified with primers: 5’-AATAGTCGACGATGAGTAAAGGAGAAGAACTTTT-3’, 5’-TATTGGGCCCTCATTACTTATTAAAAACTGGC-3’. These fragments were cloned on the sides of *neo5* immediately followed by a *MTT1* promotor to make pNeo5_ovGFP-LF4A. The pNeo5_ovGFP-LF4A plasmid was used for generating a germline integrant in which the MTT1 gene promoter is placed in front of a coding region expressing GFP-LF4A (in the *LF4A* locus) and the derived heterokaryon was used in the suppressor screen (see below).

Our *neo5* was used to make plasmids for somatic disruptions of *LF4A* and overproduction of GFP-LF4A (pNeo5_ovGFP-LF4A) in the background of MTT1-GFP-DYF1 placed in the *BTU1* locus as described [119].

To overexpress mCherry-LF4A, the GFP coding region in pNeo5_ovGFP-LF4A was replaced with that of mCherry [120], which was amplified with primers: 5’-CTAAACTTAAAATAATGGCCAAGTCGACGGTTTCAAAAGGAGAAGAAG-3’, 5’-GATAACAATTTATATTAGTTCATGACGCGTTTGTAAAGTTCATCCATACC-3’ from pNeo4-mCherry. To overexpress mCherry-LF4A^F82A^, the GFP-LF4A part of pNeo5_ovGFP-LF4A was replaced with two fragments that provide the sequence of mCherry-LF4A^F82A^ using NEBuilder Hifi DNA Assembly. The point mutation was created at the junction between the two fragments, which were amplified with the following primer pairs: 5’-CTAAACTTAAAATAATGGCCAAGTCGACGGTTTCAAAAGGAGAAGAAG-3’, 5-CAATTCAGCCACTAGTGCCAAACGTCCTGTAG-3’, 5’-GCACTAGTGGCTGAATTGATGGATCAGAACC-3’ AND 5’-CAAAAGCTGGGTACCGGGCCCATATGGGTGGCGTG-3’ from pNeo5_ovmCherry-LF4.

The pNeo5_ovmCherry-LF4A and pNeo5_ovmCherry-LF4A-F82A plasmids were also used for overexpression in the background of either MTT1_p_-GFP-DYF1 placed in the *BTU1* locus as described [119] or IFT140-GFP in its own locus using a somatic (macronuclear) approach.

### Somatic and germline transformation

All procedures for generating somatic and germline transformants and crosses were done as described [112, 113] with some changes. For the germline biolistic transformation, 100 µg of plasmid DNA was digested with restriction enzymes to release the targeting fragment, and used to coat gold particles (SeaShell S550d or Chempur 900040), which were then deposited onto seven microcarriers and fitted into the hepta adapter (Biorad). A biolistic shooting was performed 4 hours after mixing of CU428 and B2086 strains. The hepta adapter assembly was positioned on the third slot from the top inside PDS1000/He and the rapture disk had the value of 1800 psi. The conjugating cells were spread as a thin layer onto a 10-cm wide Petri dish containing a layer of Tris-Agar (10 mM Tris-HCl pH 7.4, 1.5% agar) and positioned at the bottom-most slot of PDS1000/He. To generate homokaryons, we either used the standard procedure based on two rounds of genomic exclusion or the short circuit genomic exclusion (SCGE) [121] with either B*VI or B*VII [83]. Specifically, the homokaryons for SUP1, CDKR1-KO, LF4A-KO_CDKR1-KO, ovGFP-LF4A and ovGFP-LF4A_CDKR1-KO were made using the SCGE. For genotyping the *cdkr1*^*sup1*^ allele its sequence introduced an MboI restriction site that was detected in the 1 kb PCR product (with primers 5’-TGGTGATTTTGGATCAGCT-3’ and 5’-CTTGCTTTCCTCAAATAAAC-3’).

### Confocal and TIRF microscopy

For immunofluorescence, cells were stained as described [119] using a simultaneous fixation (2% paraformaldehyde) and permeabilization (0.5% Triton X-100). To test for association with the cytoskeleton, cells were permeabilized with 0.5% Triton X-100 prior to fixation with paraformaldehyde. The primary antibodies were: anti-GFP (1:100 Abcam ab6556), anti-centrin 20H5 (1:200, Millipore 04-1624), and anti-polyglycine serum 2302 (1:200, gift of Martin Gorovsky, University of Rochester). The secondary antibodies were purchased from Jackson Immuno Research. Images were taken using a Zeiss LSM 710 or LSM 880 confocal microscope (63× oil immersion, 1× or 1.5× digital zoom) and analyzed with Fiji-ImageJ software [122]. The total internal reflection fluorescence microscopy of *Tetrahymena* was done as described [76]. Images and video recordings were processed in Fiji-ImageJ [122].

### Western blots

Protein samples were separated on a 10% SDS-PAGE. The gels were either stained with blue silver [123], or proteins were transferred onto a PVDF membrane for western blotting. Primary antibodies used with western blots were: anti-GFP (Abcam ab6556 at 1:5000 or Rockland 600-401-215 at 1:1000), and the anti-thiophosphate ester 51-8 (1:2000-5000, Abcam ab92570). Colorimetric images of stained gels and luminescent images of western blots were recorded using ChemiDoc MP System and processed with Image Lab (Biorad).

### *In vitro* kinase assay

*Tetrahymena* cultures at 2×10^5^ /ml (30 ml) of strains expressing GFP-LF4A were treated with CdCl_2_ (2.5 µg/ml) for 3-5 hours, cells were collected at 1700 rcf for 3 min and washed once with 10 mM Tris-HCl (pH 7.5). A lysis buffer (5 ml per 30 ml of Cd^2+^ induced cells) was added that contained 0.5% NP-40, a phosphatase inhibitor cocktail (20 mM beta-glycerophosphoate, 1 mM sodium orthovanadate, 1 mM Na4P2O7, 20 mM NaF) and 70 mM E64, in TBS (20 mM Tris-HCl pH 7.5, 150 mM NaCl). PMSF was added to 1 mM immediately before suspending the cell pellets. The mixtures were kept on ice and were pipetted vigorously every 10 minutes for 3 times. Then, the mixtures were spun at 20,000 rcf for 15 minutes. The supernatants were collected, diluted with 2 volumes of TBS so that the NP-40 concentration is below 0.2% for immunoprecipitation with GFP-Trap beads (agarose, ACC0CM-GFA0050, Allele). The GFP-Trap beads were added to the diluted lysate (10-15 µl resin per 30 ml of the starting culture). The mixtures were then incubated at 4 °C for 2 hr. The beads were pelleted at 2500 rcf for 2 min, washed with: TBS, TBS with 500 mM NaCl, and the kinase buffer (50 mM Tris-HCl pH 7.2, 100 mM NaCl, 10mM MgCl_2_). A total of 30 µl of the kinase buffer with 1 mM DTT, 100 μM ATP-gamma-S (Biolog 88453-52-5) and phosphatase inhibitors cocktails were added to no more than 15 µl of the beads slurry along with 20 µg MBP (Active Motif 31314). The kinase assay mixtures were incubated for 30-120 minutes at 30°C. The reactions were either stored overnight refrigerated or immediately terminated by addition of 2.5 mM PNBM (Abcam ab138910) followed by incubation for 2 hours at room temperature. The samples were then treated with the sample buffer and separated by SDS-PAGE. The thio-phosphorylation was detected using anti-thiophosphate ester 51-8 (1:2000-5000, rabbit, Abcam ab92570).

### Genetic screen for suppressors of GFP-LF4A overexpression

The neo5-MTT1_p_-GFP-LF4a fragment was targeted to the *LF4A* locus in the micronucleus by biolistic transformation as described [112] using the hepta adapter as outlined above. The heterokaryon strain named ovGFP-LF4A-HE-3B (mic: neo5-ovGFP-lf4a/neo5-ovGFP-lf4a mac: wildtype, VII) was grown to 1×10^5^ cells/ml (40 ml total), and treated 10 µg/ml of nitrosoguanidine (Sigma-Aldrich) for 3 hours at 30°C. The mutagenized culture was subjected to starvation in Dryl’s buffer [124](1.7 mM sodium citrate, 1 mM NaH_2_P0_4_, 1 mM Na_2_HPO_4_ and 1.5 mM CaCl_2_), overnight at 30°C. The mutagenized cells were mixed with an equal volume and number of starved B*VI and subjected to uniparental cytogamy [79]. Briefly, 5-6 hours after strain mixing, the conjugating cells were treated with 1.4% glucose for 45 minutes (by addition of an appropriate volume of 20% glucose) and diluted with 7-8 volumes of sterile water. The cells were spun down, suspended in 30 ml of SPPA and split into 5 ml samples, each kept in a 500 ml round media bottle (six total) at room temperature overnight to complete conjugation. SPPA (45 ml) was added to each bottle, followed by incubation at 30°C for 3 hr. Paromomycin was added to the final concentration of 200 µg/ml to select the conjugation progeny. After 24-36 hours of selection at 30°C, the pm-r cells were collected by centrifugation, each sample was suspended in fresh 40 ml SPPA with 2.5 µg/ml CdCl_2_ and incubated in 50 ml conical centrifuge tubes in vertical positions overnight at room temperature. Due to the GFP-LF4A overproduction and the resulting loss of cilia, unsuppressed cells become paralyzed and sunk to the bottom of the tubes. Cells carrying suppressor mutations remained motile neat the tube top. The supernatants (4-5 ml) were collected from the top of each vertically positioned tube and the cells were transferred into 5 ml fresh SPPA in 15 ml conical tubes, incubated horizontally for 6 hours to overnight, then washed and suspended in 10 ml SPPA with 2.5 µg/ml CdCl_2_ and the tubes were kept vertically oriented overnight. Single clones were isolated from the top 1 ml of culture of each tube and retested for suppression on 96-well plates. The F0s (isolated suppressor clones) were matured and mated to the cycloheximide resistance (cy-r) heterokaryon CU427 (mic: *chx1-1*/*chx1-1*, mac: + mt VI) or CU427.7 (mic: *chx1-1*/*chx1-1*, mac: + mt VII) and the F1 progeny was recovered as pm-r and cy-r cells. The F1s were matured and allowed to assort to paromomycin sensitivity (pm-s). To make F2s, assorted pm-s F1 were subjected to SCGE with B*VI or B*VII as described [83]. The F2s were cloned by picking pm-r cells from independently selected wells, grown in SPPA, and replica-plated on SPPA with 2.5 µg/ml CdCl_2_ to test for paralysis.

### Identification of the causal mutations for suppressors

For the intragenic suppressors, total genomic DNA was extracted from a pool of F2s and the several 1-1.2 kb overlapping fragments covering MTT1p-ovGFP-LF4A transgene were amplified and sequenced using the Sanger method.

Pools of clones were prepared containing pm-r F2s obtained by SCGE from a single sup1/sup1^+^ F1. The unsuppressed pool contained 12 F2s clones that consistently became paralyzed in Cd^2+^ and the suppressed pool contained 14 F2 clones that remained motile in Cd^2+^. The two pools were grown in 25 ml volumes, starved for 2 days at room temperature in 60 mM Tris-HCl and the total genomic DNAs of the pools were extracted, using the urea method [125]. The genomic DNAs were subjected to whole genome sequencing using Illumina technology exactly as described [83]. Sequences were aligned to the macronuclear reference genome (June 2014 version, GenBank assembly accession GCA_000189635.1) [126] and variants were detected filtered and subtracted as described [83]. In parallel, the suppressor and non-suppressor reads where aligned to the micronuclear reference genome [127] and the allelic contrast analysis was performed as described [83] to detect a micronuclear chromosome location with increased frequency of variant cosegregation with the mutant (suppressed) phenotype using MiModD (Version 0.1.8 [128]).

### Structural modeling of LF4A and CDKR1 kinases

Curated multiple sequence alignment profiles of protein kinases from diverse organisms were used to classify CDKR1 [68, 129, 130]. MAPGAPS [131] and HMMER [132] were used to detect and align CDKR1 to the best matching CDC2 profile [133].

The template for 3D homology modeling of LF4A was identified by performing a BLAST search against the PDB database using blastp routine from the NCBI [134]. The cyclin-dependent kinase from *Cryptosporidium* (Chain A of PDB 3NIZ, 37% sequence identity) was chosen as the template. MODELLER (version 9.12) was used to generate the homology model for LF4A from 3NIZ using the automodel module [81]. ADP and magnesium ions present in the template were also included in the modeled structure of LF4A. Jpred [88] was used to generate a secondary structure prediction of the CDKR1 (TTHERM_01080590) sequence. To make a structural model of CDKR1, *ab initio* protein structure prediction algorithms and I-TASSER were used [135-138]. The visualization of the modeled structures was performed using PyMOL [135].

## Acknowledgements

This work was supported by the NIH grants R21HD092809 (to JG) and 5RO1GM114409 (to NK), a bridge funding from the Office of the Vice-President for Research and the Department of Cellular Biology at the University of Georgia (to JG), the National Science Foundation grant MCB-1149106 (to NK), grants from Deutsche Forschungsgemeinschaft BIOSS CRC746 and CRC850 (to RB), and grants from the National Science Centre, Poland: OPUS13 2017/25/B/NZ3/01609 (to DW) and OPUS15 2018/29/B/NZ3/02443 (to EW). GM was supported by a pre-doctoral fellowship from the National Institutes of Health (1F31NS074841-01). We thank Martin A. Gorovsky (University of Rochester) for the polyG antibodies. We also acknowledge the assistance of Muthugapatti K. Kandasamy at the UGA Biomedical Microscopy Core with confocal imaging.

## Supporting information

**S1 Figure.**
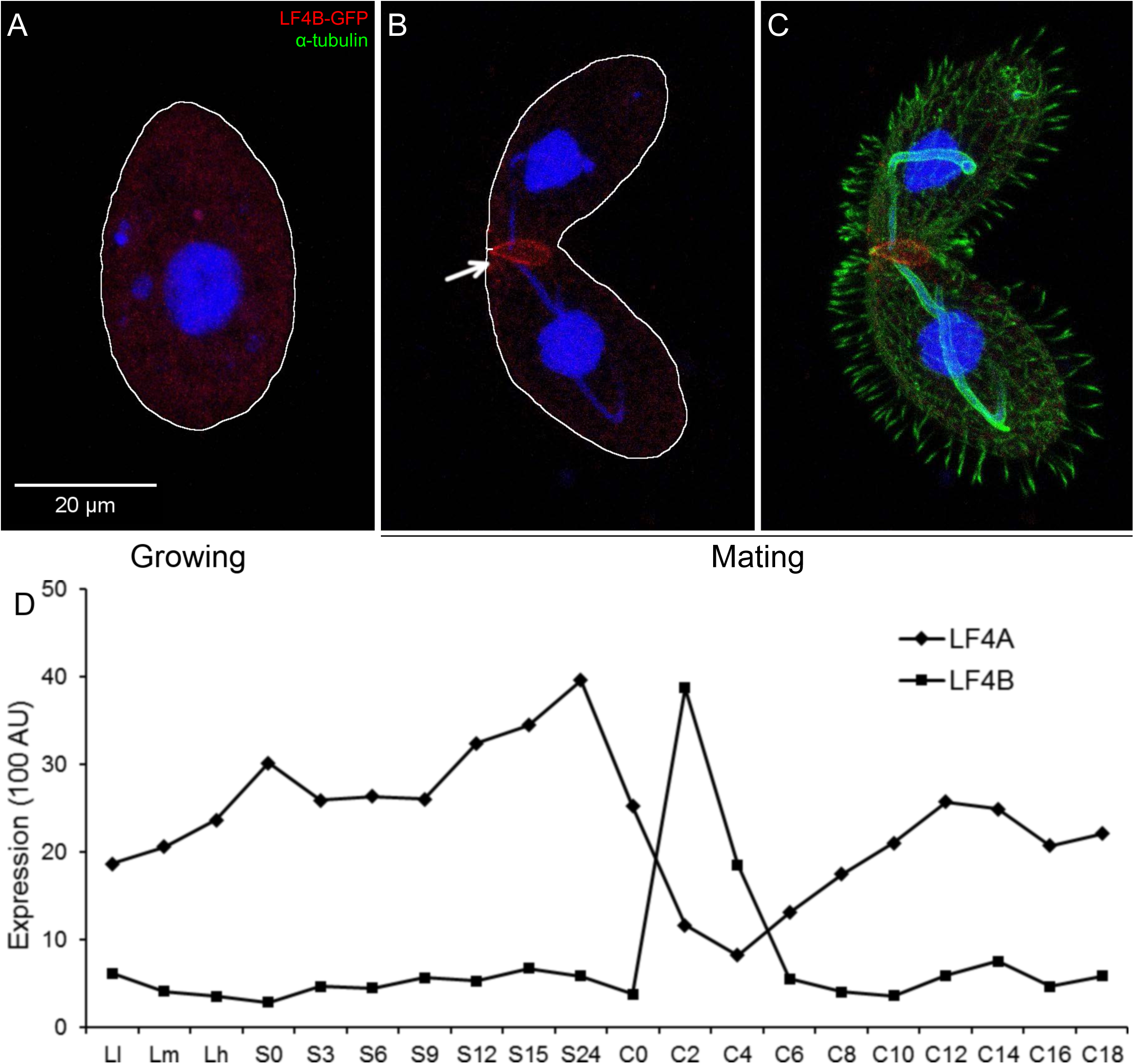
L4B is associated with the cell-cell junction during conjugation. (A-C) Cells expressing LF4B-GFP (the tag added by engineering the native locus) analyzed by immunofluorescence using anti-GFP antibodies (red) 12G10 anti-*α*-tubulin (green only in panel C) and DAPI (Blue). (A) A vegetatively growing cell. Note an absence of a GFP signal above the typical background. (B-C) A conjugating pair. Note that LF4B-GFP localizes to the junction between the two mating cells. (D) Expression profiles of mRNAs for LF4A (TTHERM_00058800) and LF4B (TTHERM_00822360) obtained from the *Tetrahymena* Functional Genomics database (http://tfgd.ihb.ac.cn/search/detail/gene/TTHERM_00822360). The levels of mRNA at the following conditions are shown: L-l, L-m and L-h: vegetatively growing cells collected at ∼1×10^5^ cells/ml, ∼3.5×10^5^cells/ml and ∼1×10^6^ cells/ml. S-0, S-3, S-6, S-9, S-12, S-15 and S-24: cells starving for 0, 3, 6, 9, 12, 15 and 24 hours. C-0, C-2, C-4, C-6, C-8, C-10, C-12, C-14, C-16 and C-18: conjugating cells collected at 0, 2, 4, 6, 8, 10, 12, 14, 16 and 18 hours after initiation of conjugation by mixing different mating types.

**S2 Figure.**
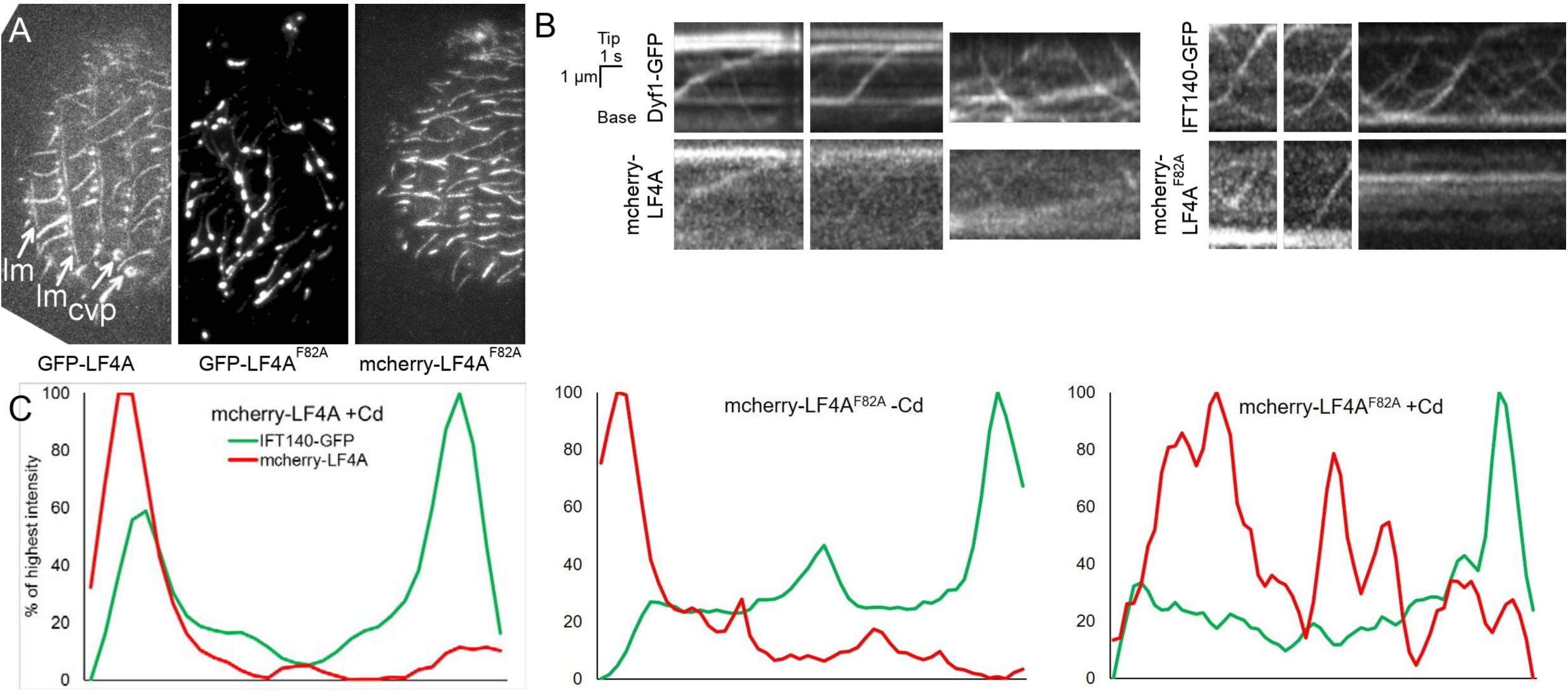
Association of GFP-LF4A with microtubules and its co-transport with IFT. (A) TIRF images of live cells overexpressing GFP-LF4A (left), kinase-weak GFP-LF4A^F82A^ variant (middle), and kinase weak mCherry-LF4A^F82A^. Overexpressed GFP-LF4A (right panel) localized to the bases of cilia and along cilia but also near the microtubule-rich structures in the cell body including longitudinal microtubules (lm), and contractile vacuole pores (cvp). The kinase-weak GFP-LF4A^F82A^ is enriched at the tips of cilia while mCherry-LF4A^F82A^ is distributed uniformly along cilia. (B) Kymographs that document co-migration of IFT proteins (GFP-DYF1 or IFT140-GFP, top) and either mCherry-LF4A or mCherry-LF4AF82A after induction with Cd^2+^ (3 hours). (C) Signal intensity profiles of single cilia in cells expressing either mCherry-LF4 or mCherry-LF4F82A (red) and IFT140-GFP (green). The base is on the left and the tip is on the right side of each profile. Note that the active kinase is enriched at the base (left profile). The weak kinase is enriched at the base and spread along the cilium length when overproduced but does not accumulate at the tip. The pattern distribution of IFT140 is similar in all background and conditions, with enrichment at the tip.

**S3 Figure.**
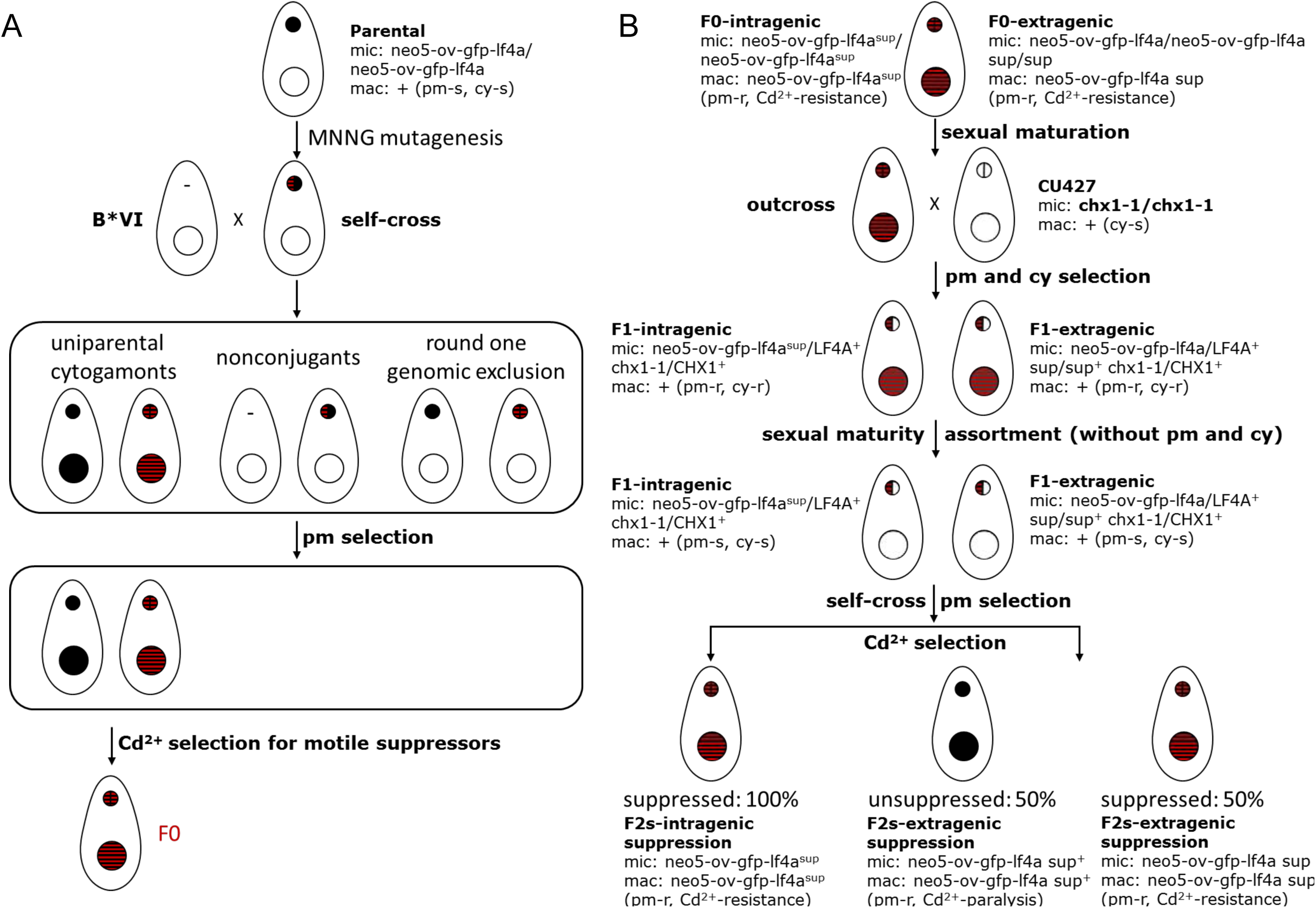
A detailed presentation of the pipeline used for identifying suppressors of overexpression of GFP-LF4A. (A) Steps involved in generation and isolation of the suppressor F0s. A heterokaryon with the ovGFP-LF4A transgene in the micronucleus (solid black) was subjected to mutagenesis with nitrosoguanidine. The mutagenized heterokaryon was subjected to a self-cross (uniparental cytogamy) that involves a mating to a star strain that lacks a functional micronucleus. The outcome includes the desired self-cross progeny (uniparental cytogamonts, typically a few % of the conjugated pairs), cells that failed to undergo conjugation (nonconjugants) and round one genomic exclusion, the most common outcome of such a cross (typically >95%). The uniparental cytogamy progeny were selected with paromomycin (pm) as they expressed the transgene in the macronucleus. The suppressor F0s were then isolated by collecting cells that remained mobile after overnight Cd^2+^ exposure. (B) Steps used to determine whether the suppression is intragenic or extragenic. Each suppressor F0 clone underwent sexual maturation and was mated to CU427, a strain with a micronucleus carrying a homozygous *chx1-1* allele (resistance to cycloheximide cy) and a wild-type macronucleus. The outcross progeny was selected with cy and pm. F1 clones underwent phenotypic assortment to pm-s and become sexually mature. The pm-s F1 clone was then subjected to self-cross (short-circuit genomic exclusion) and the pm-r F2 clones were obtained. A number of F2 clones of each suppressor were tested for suppression by Cd^2+^ treatment. An intragenic suppressor gives only suppressed F2 clones. An extragenic suppressor gives both unsuppressed (paralyzed) and suppressed F2 progeny.

**S4 Figure.**
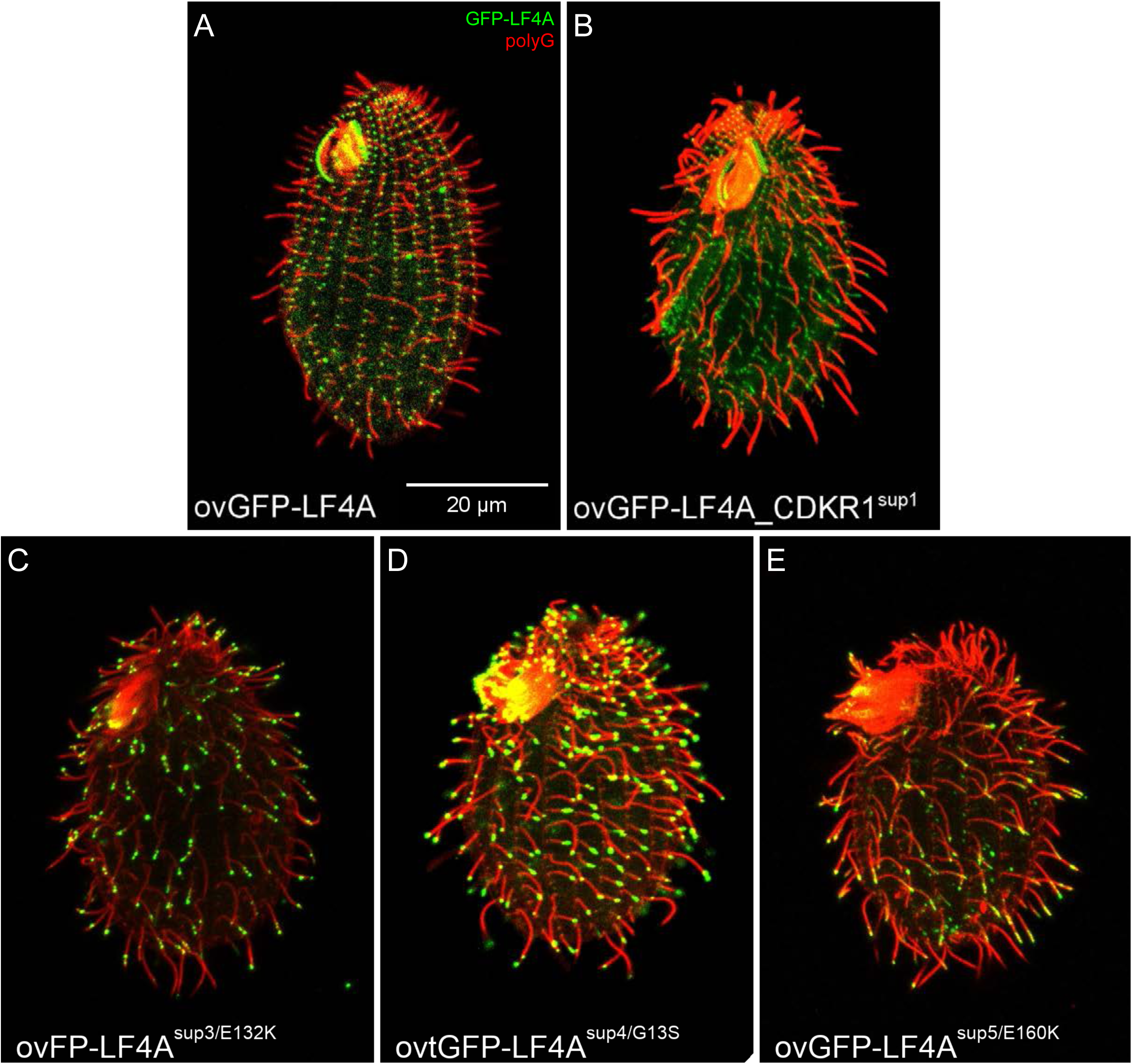
Phenotypes of intragenic and extragenic suppressors of GFP-LF4A overexpression. Self-cross progeny at a control background of GFP-LF4A overexpression (A), the extragenic suppressor SUP1 (B) and three intragenic suppressors SUP3, SUP4 and SUP5 (C-E). All cells were subjected to a 6-hours Cd^2+^ exposure prior to immunofluorescence assay and showed the GFP signal (green) and were stained with anti-polyG antibodies (red).

**S5 Figure.**
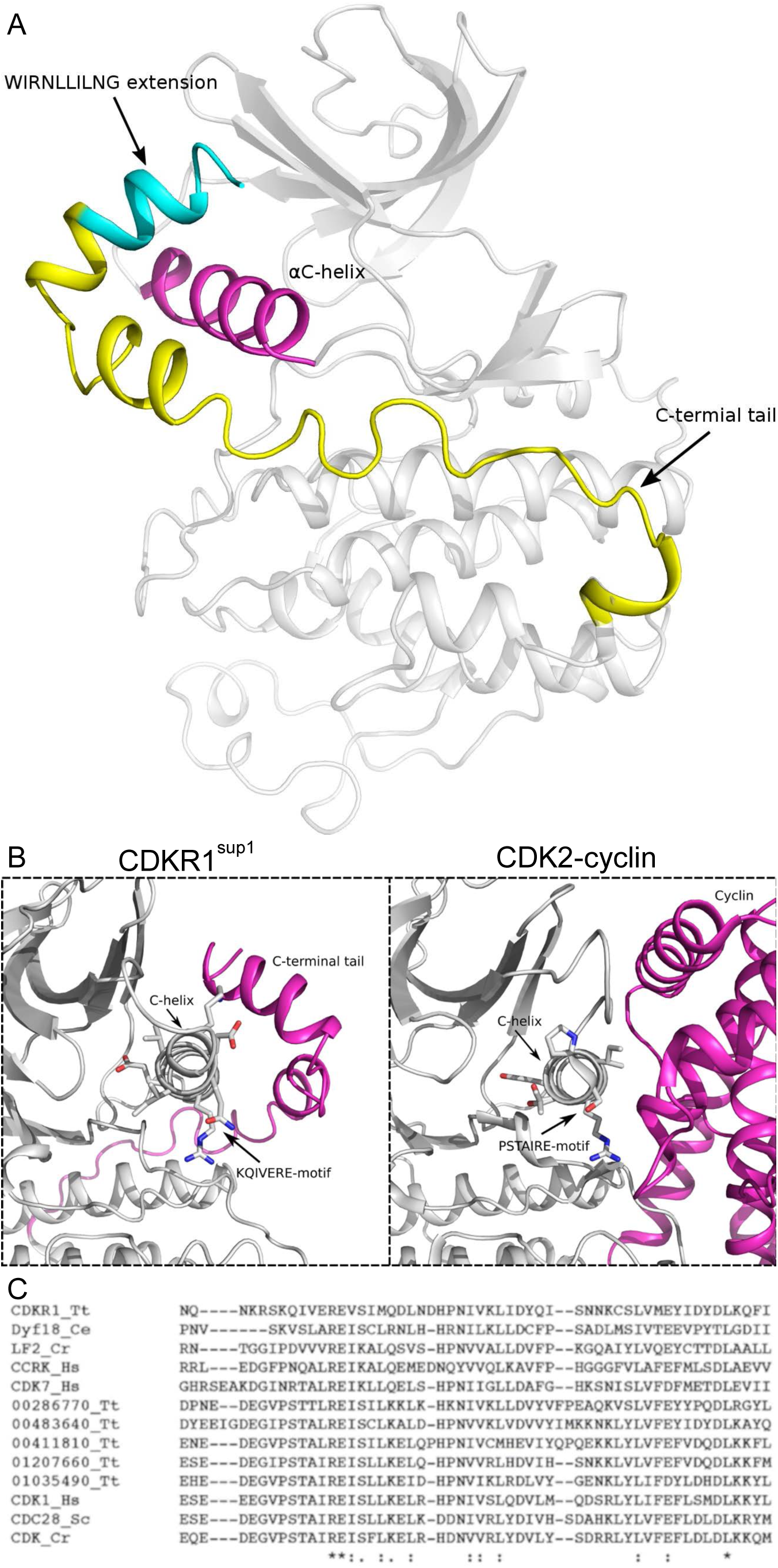
A predicted structure suggests that the C-terminal tail extension in CDKR1^sup1^ affects the kinase function. (A) Predicted structure of CDKR1^sup1^ with a C-terminal tail (with a WIRNLLILNG extension) forming two helical segments on the top of C-helix. (B) 3D view comparison of the C-helix (and cyclin-CDK interface) of CDKR1^sup1^ and a CDK2. A PSTAIRE sequence in the canonical CDKs lies at the interface of the cyclin-CDK complex and corresponds to the C-helix in the CDKs. The equivalent positions in CDKR1 are KQIVERE. (C) A sequence alignment of fragments of CDK and CDK-related kinases.

**S6 Figure.**
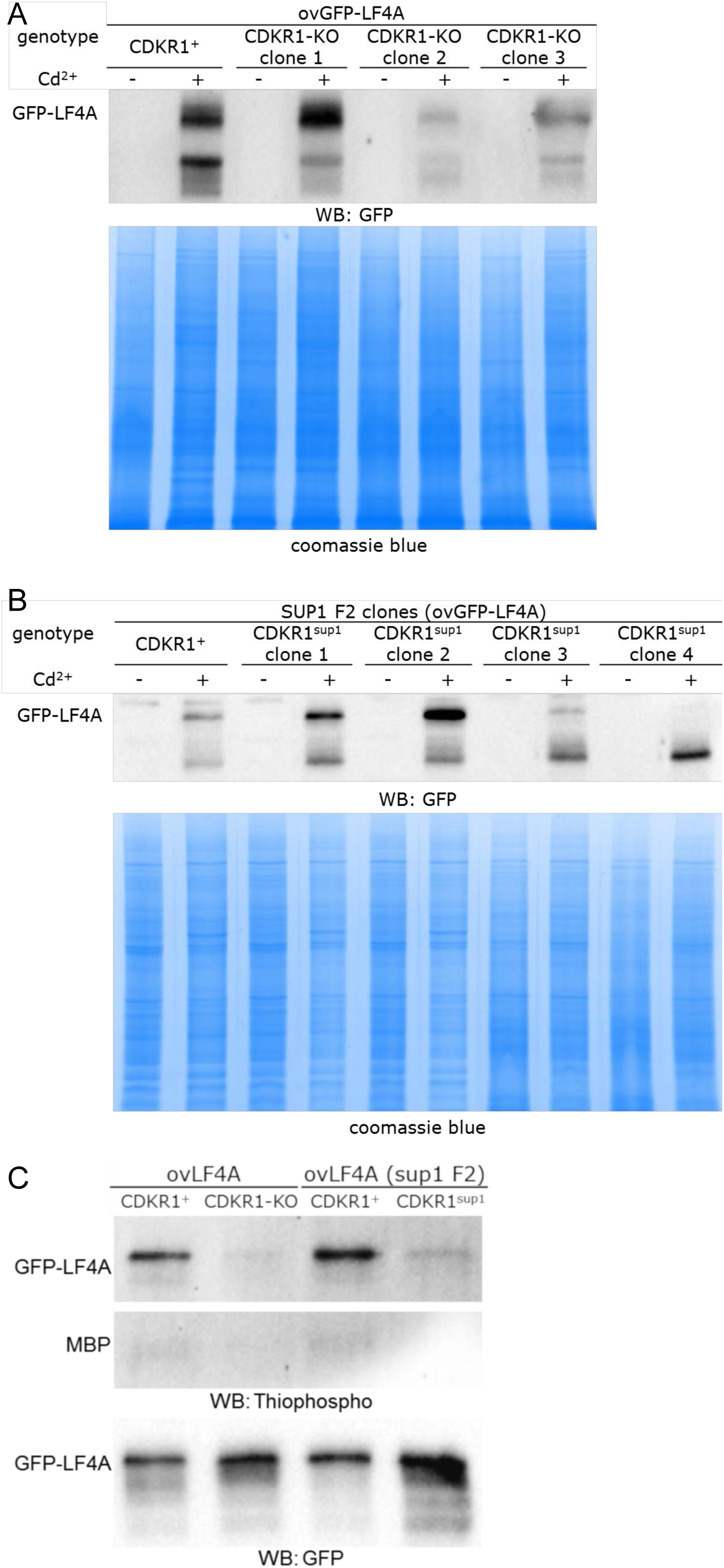
A loss of CDKR1 does not consistently affects the levels of overproduced GFP-LF4A. (A) A comparison of the levels of GFP-LF4A in whole cell lysates of several F2 clones derived from the same F1, with or without a 6-hour Cd^2+^ exposure. An ovGFP-LF4A_CDKR1^+^ control strain and three ovGFP-LF4A_CDKR1-KO F2s were analyzed. (B) A comparison of the levels of GFP-LF4A in whole cell lysates of the F2 progeny clones of the extragenic suppressor SUP1, with or without a 6-hour Cd^2+^ exposure. An unsuppressed (ovGFP-LF4A_CDKR1^+^) strain and four suppressed (ovGFP-LF4A_CDKR1^sup1^) strains were analyzed. Faster-migrating bands represent degradation products of GFP-LF4A. (C) In vitro kinase assays show that loss-of-function of CDKR1 result in a reduced kinase activity of overproduced GFP-LF4A against itself and MBP. The top panel is a western blot that reveals the signal of thiophosphorylated substrates. The bottom panel is a western blot that documents the levels of GFP-LF4A in the reactions using anti-GFP antibodies

**S1 Video.** A live cell expressing LF4A-GFP (under the native promoter) recorded by TIRFM. The frame rate is 3 times of the real time. Examples of rare cilia with mobile LF4A-GFP are marked by red boxes.

**S2 Video.** A TIRFM video of a live wild-type cell expressing GFP-DYF1 reporter (treated with Cd2+ for 3.5 hours). The frame rate is 3 times of the real time.

**S3 Video.** A TIRFM video of a live cell that overexpresses GFP-LF4A (treated with Cd2+ for 3.5 hours) that also expresses GFP-DYF1 reporter. The frame rate is 3 times of the real time.

**Table S1** The candidate causal variants identified in the SUP1 suppressor genome based on bioinformatic subtractions and filtering.

